# Selective Impact of ALK and MELK Inhibition on ERα Stability and Cell Proliferation in Cell Lines Representing Distinct Molecular Phenotypes of Breast Cancer

**DOI:** 10.1101/2023.12.19.572304

**Authors:** Stefania Bartoloni, Sara Pescatori, Fabrizio Bianchi, Manuela Cipolletti, Filippo Acconcia

## Abstract

Breast cancer (BC) is a leading cause of global cancer-related mortality in women, necessitating accurate tumor classification for timely intervention. Molecular and histological factors, including PAM50 classification, estrogen receptor α (ERα), breast cancer type 1 susceptibility protein (BRCA1), progesterone receptor (PR), and HER2 expression, contribute to intricate BC subtyping. Through in silico screenings and multiple BC cell line investigations, we identified enhanced sensitivity of ERα-positive BC cell lines to ALK and MELK inhibitors, inducing ERα degradation and diminishing proliferation in specific BC subtypes. MELK inhibition attenuated ERα transcriptional activity, impeding E2-induced gene expression, and hampering proliferation in MCF-7 cells. Synergies between MELK inhibition with 4OH-tamoxifen (Tam) and ALK inhibition with HER2 inhibitors revealed potential therapeutic avenues for ERα-positive/PR-positive/HER2-negative and ERα-positive/PR-negative/HER2-positive tumors, respectively. Our findings propose MELK as a promising target for ERα-positive/PR-positive/HER2-negative BC and highlight ALK as a potential focus for ERα-positive/PR-negative/HER2-positive BC. The synergistic anti-proliferative effects of MELK with Tam and ALK with HER2 inhibitors underscore kinase inhibitors’ potential for selective treatment in diverse BC subtypes, paving the way for personalized and effective therapeutic strategies in BC management.

## Introduction

Breast cancer (BC) remains the most lethal neoplastic disease affecting women worldwide. Early diagnosis requires the accurate classification of mammary tumors to determine the appropriate pharmacological approach, based on various criteria. The classification of breast tumors involves the molecular expression of specific genes using the PAM50 classification, which categorizes them into five clinicopathological surrogates: luminal A (LumA), luminal B (LumB), HER2-overexpressing (HER2+), basal epithelial-like (BL), and normal-like (NL). Additionally, the histological type of the tumor (e.g., invasive ductal carcinoma - IDC, adenocarcinoma, papillary carcinoma) is an important tool for characterizing mammary carcinomas. Several key prognostic factors for BC include the expression of estrogen receptor α (ERα), which distinguishes tumors as ERα-positive or ERα- negative, the status of breast cancer type 1 susceptibility protein (BRCA1) (wild type – wt versus mutated), and the expression of progesterone receptor (PR) and HER2, further dividing different subgroups within the LumA and LumB phenotypes. However, there is some overlap between tumor classifications, as any histological tumor type can be both ERα-positive and ERα-negative and belong to different clinical surrogates of BC. For example, LumA tumors (PR-positive/HER2-negative; PR-negative/HER2-negative) and LumB tumors (PR-negative/HER2-positive; PR-positive/HER2-positive) are ERα-positive, while the other subtypes are ERα-negative, and BRCA1-mutated carcinomas do not express ERα, but all of them can originate from various histological types ^1–4^. Therefore, BC includes a variety of different molecular and biological phenotypes that make it a jumble of single intrinsically different diseases.

Upon diagnosis, approximately 70% of newly detected breast tumors express ERα and exhibit a more favorable prognosis compared to ERα-negative tumors. This is attributed to the fact that ERα serves as the pharmacological target for ERα-positive tumors, which are treated with endocrine therapy drugs that hinder various aspects of the 17β-estradiol (E2):ERα signaling pathway to impede cell proliferation. Patients are prescribed either aromatase inhibitors (AIs) to suppress E2 production, selective estrogen receptor modulators (SERMs) like 4OH-Tamoxifen (Tam) to inhibit ERα transcriptional activity, or selective estrogen receptor down-regulators (SERDs) such as fulvestrant to induce ERα 26S proteasome-dependent degradation ^1–4^. However, LumA and LumB tumors show different sensitivities to ET drugs. Tam is the primary clinical treatment for LumA tumors, whereas LumB tumors, which express HER2, necessitate combination therapy involving Tam along with drugs targeting this additional molecular target (e.g., gefitinib - Gef, lapatinib - Lapa, and erlotinib - Erlo) ^4,5^. Therefore, the correct classification of the mammary tumor determines the specific pharmacological approach for patients.

Despite the established effectiveness, ongoing treatment of patients with ET results in the development of drug-resistant tumors in approximately 50% of cases, leading to relapse and metastatic recurrence in distant organs. Metastatic breast cancer (MBC) cells, which retain ERα expression, do not respond to ET drugs and prove exceedingly challenging to treat, often resulting in a fatal outcome. Furthermore, different subtypes of MBC exist, representing distinct diseases and contributing to the increased variability of overall BC phenotypes ^1–5^.

The substantial heterogeneity of BC and MBC phenotypes, coupled with the development of resistance to ET drugs, underscores the need to identify novel therapeutics that selectively target specific BC subtypes. Such drugs would either prevent the emergence of drug resistance or effectively combat metastatic disease. Recently, our research has demonstrated that drugs capable of inducing ERα degradation through diverse mechanisms inhibit BC cell proliferation. This finding has allowed us to identify several Food and Drug Administration (FDA)-approved drugs, initially designed for different purposes, which possess ‘anti-estrogen-like’ properties, inducing ERα degradation and effectively halting the proliferation of BC cell lines ^6–14^.

Interestingly, among the identified drugs, we found that the anti-proliferative effects of cardiac glycosides (CG) ouabain and digoxin are more pronounced in ERα-positive BC cell lines compared to ERα-negative ones, primarily due to their ability to induce ERα degradation ^6,10^. These findings suggest that ERα-positive breast tumor cells might exhibit higher sensitivity to specific drugs compared to ERα-negative breast tumor cells, as these drugs induce the degradation of ERα, a transcription factor crucial for the G1 to S phase progression of the cell cycle ^15^. Additionally, we made an unexpected discovery that the GART inhibitor lometrexol is effective only in LumA IDC cells, which mimic both primary and metastatic BC ^16^, while the CHK1 inhibitors AZD7762 and prexasertib lead to ERα degradation and prevent the proliferation of cell lines mimicking the LumA but not the LumB tumor phenotype ^17^. Therefore, drugs inducing ERα degradation can specifically reduce the proliferation of certain BC subtypes.

This evidence suggests the existence of drugs inducing ERα degradation that could exhibit enhanced sensitivity in ERα-positive compared to ERα-negative breast tumor cells and could selectively target specific subtypes of ERα-positive BC. To explore this hypothesis, we conducted experimental investigations utilizing a combination of screenings in silico and across various BC cell lines. Our findings revealed that the inhibition of anaplastic lymphoma kinase (ALK) and maternal embryonic leucine zipper kinase (MELK) selectively induces ERα degradation and prevents the proliferation of cell lines representing the LumB ERα-positive/PR-negative/HER2-positive and LumA ERα-positive/PR-positive/HER2-negative molecular phenotypes of BC, respectively.

## Results

### Identification of ALK as a kinase regulating ERα stability

We employed an unbiased approach to identify drugs with increased sensitivity in ERα-positive breast cancer (BC) cell lines compared to ERα-negative ones. Our investigation involved analyzing the DepMap portal (https://depmap.org/portal/), which contains data on approximately 4600 drugs and their effects on the cell proliferation of 26 BC cell lines. Each drug’s sensitivity in specific BC cell lines is represented by a numerical value in the DepMap portal. To stratify the BC cell lines based on ERα expression, we utilized previous molecular characterizations of the BC cell lines ^18,19^. For each drug, we calculated the mean sensitivity value in both ERα-positive and ERα-negative BC cell lines. Then, we determined the difference in mean sensitivity between ERα-positive and ERα-negative BC cell lines for each drug. Using a Student t-test, we estimated the relative p-values, which were subsequently -Log_2_ transformed. We visualized the results in a Volcano plot, revealing that the majority of drugs in the DepMap database exhibited increased sensitivity in ERα-positive BC cell lines compared to ERα-negative ones (Fig. 1A and Supplementary Table 1).

**Figure 1.**
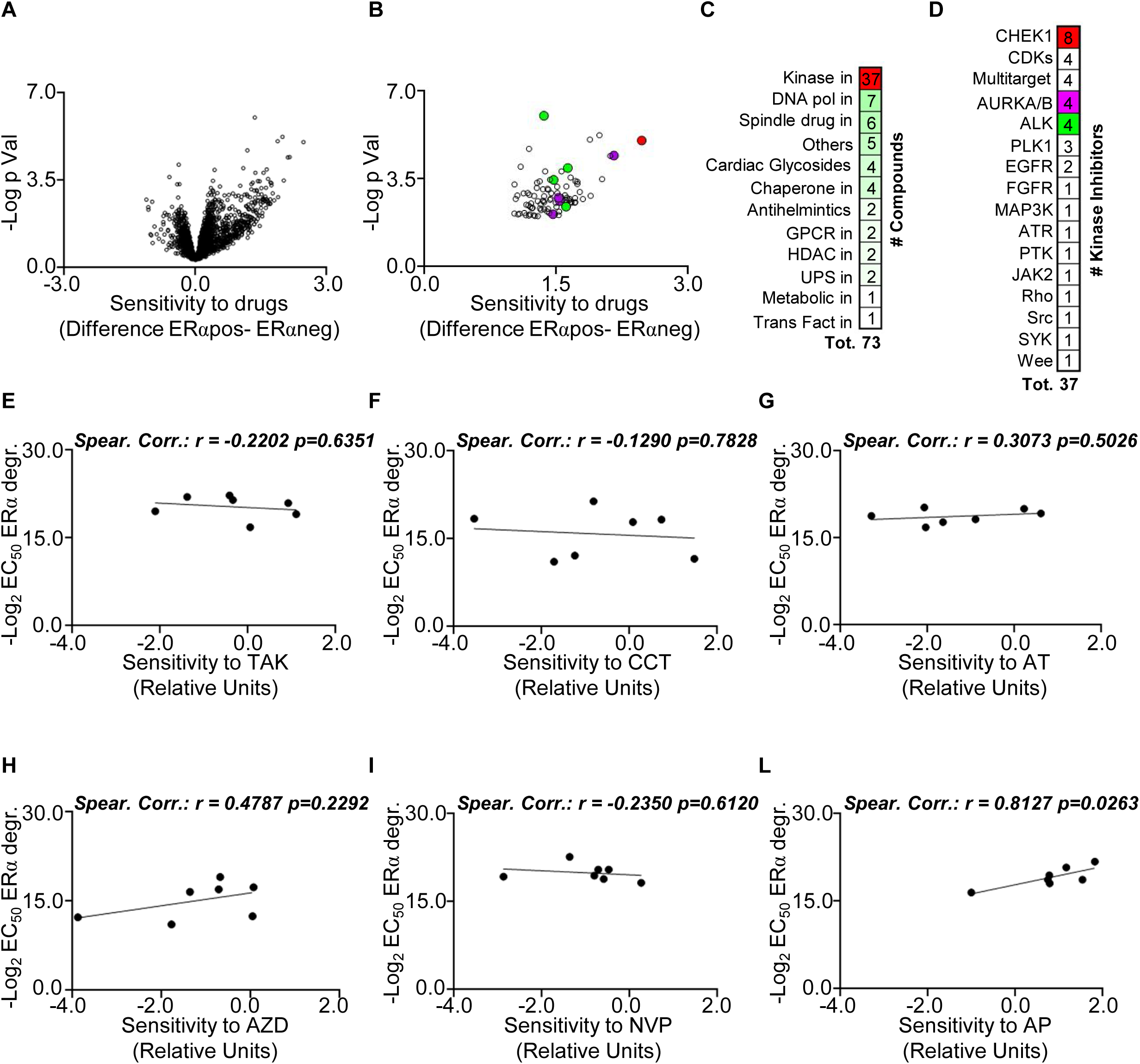
Potential Regulation of ERα Stability by ALK Kinase. (A) Volcano plot illustrating differences in drug sensitivity between ERα-positive and ERα-negative breast cancer (BC) cell lines. Data sourced from the DepMap portal (https://depmap.org/portal). Each dot represents a drug’s value in the database. (B) Volcano plot revealing differences in drug sensitivity between ERα-positive and ERα-negative BC cell lines, after applying the specified thresholds (please see the text) for positive hit selection. Each dot represents a drug’s value in the database, and color dots correspond to drugs highlighted in panels C and D. (C) Number of compounds identified as positive hits in panel (B), categorized as indicated alongside panel C. (D) Number of kinase inhibitors identified as positive hits in panel (C), with the target of each kinase inhibitor specified alongside panel D. (E-F) Linear regression and Spearman Correlation values between the sensitivity to AURKA/AURKB inhibitors TAK901 – TAK (E), CCT137690 – CCT (F), AT9283 – AT (G), or to ALK inhibitors AZD3436 – AZD (H), NVP-TAE684 – NVP (I), and AP26113 – AP (L), as downloaded from the DepMap portal (https://depmap.org/portal), and the effective concentration 50 (EC_50_) for inhibitor-induced reduction in ERα intracellular levels in corresponding BC cell lines. Main panels display the correlation coefficient (r) and p-values.

To identify drugs that more likely preferentially affect ERα-positive BC cell lines, we applied specific thresholds. We selected drugs with a difference in mean sensitivity between ERα-positive and ERα-negative BC cell lines greater than 1 and a corresponding p-value < 0.01. By applying these criteria, we compiled a list of 73 drugs (Fig. 1B and Supplementary Table 1). Notably, this list included cardiac glycosides (CG) and anti-helminthic drugs, known to induce ERα degradation in BC cells ^6,7,10^, as well as drugs targeting DNA polymerase or the spindle, which can induce replication stress ^20^ and potentially reduce receptor expression in BC cells ^17^. Additionally, inhibitors of the ubiquitin proteasome system (UPS), known to affect ERα stability ^11^, were present in the list (Fig. 1C and Supplementary Table 1).

A significant portion (37 drugs) of the drugs displaying increased sensitivity in ERα-positive BC cell lines compared to ERα-negative ones were kinase inhibitors (Fig. 1C and Supplementary Table 1). Among these, CHK1 was the most targeted kinase (8 drugs), and 1 inhibitor targeted ATR. Interestingly, previous research has shown that inhibiting the ATR/CHK1 pathway induces replication stress-dependent ERα degradation ^17^. Additionally, a PLK1 inhibitor was also observed in the list, and inhibition of PLK1 was previously reported to induce ERα degradation in BC cells ^21,22^. Notably, the most represented or highest-valued kinase inhibitors in terms of mean difference sensitivity among ERα-positive and ERα-negative BC cell lines or p-value were those targeting ALK or AURKA and AURKB (Fig. 1D and Supplementary Table 1).

In light of these findings, we proceeded to examine the effects of three inhibitors of ALK (namely AZD3436 – AZD, NVP-TAE684 – NVP, AP26113 – AP) and AURKA/AURKB (TAK901 – TAK, CCT137690 – CCT, AT9283 – AT) to determine their capacity to induce a reduction in ERα levels. For screening purposes the experiments were repeated twice and generated dose-response curves in seven ERα-positive BC cell lines that represent diverse clinical surrogates, histological types, and variations in PR and HER2 expression (MCF-7, ZR-75-1, T47D-1, HCC1928, BT-474, MDA-MB-361, and EFM192C cells) (Table 1) ^18,19^. Subsequently, we derived the effective dose 50 (ED_50_) for the reduction in receptor levels, which we logarithmically transformed (-Log_2_) to gauge the sensitivity of each cell line to each kinase inhibitor. We then compared these sensitivity values with the corresponding cell proliferation sensitivity values obtained from the DepMap portal for each cell line. Utilizing linear regression analyses, we found no significant correlation between any of the AURKA and AURKB inhibitors (Fig. 1E-1G and Supplementary Table 2). However, a noteworthy linear correlation (r=0.8127; p=0.0263) was observed when cells were treated solely with the ALK inhibitor AP (Fig. 1H-1L; Supplementary Table 2). Furthermore, we observed a linear correlation (r=0.7941; p=0.0329) between the sensitivity to ERα degradation in the seven cell lines for two out of three ALK inhibitors (AZD and AP) (Supplementary Fig. 1 and Supplementary Table 2). These results prompted us to conduct further investigations to validate the impact of ALK on the regulation of both ERα levels and cell proliferation.

**Table 1:** Different classification of the breast cancer cell lines used. ERα: estrogen receptor α; PR: progesterone receptor; HER2: Human Epidermal Growth Factor Receptor 2; IDC: invasive ductal carcinoma; LumA: luminal A; LumB: luminal B.

| *Cells* | *ERα* | *PR* | *HER2* | *Histotype* | *PAM50* |
| --- | --- | --- | --- | --- | --- |
| *MCF-7* | + | + | - | IDC | LumA |
| *T47D-1* | + | + | - | IDC | LumA |
| *ZR-75-1* | + | - | - | IDC | LumA |
| *HCC1428* | + | + | - | Adenocarcinoma | LumA |
| *BT-474* | + | + | + | IDC | LumB |
| *MDAMB361* | + | - | + | Adenocarcinoma | LumB |
| *EFM192C* | + | + | + | Adenocarcinoma | LumB |

### Identification of MELK as a kinase regulating ERα stability

Recently, we demonstrated that the antiviral drug telaprevir (Tel) induces degradation of the ERα and hampers the proliferation of several ERα-positive BC cell lines ^23,24^. Given the sensitivity of ERα-positive BC cell lines to various kinase inhibitors (Fig. 1), and the fact that we previously discovered that Tel inhibits the IGF1-R and AKT kinases in BC cells by reducing their intracellular levels and phosphorylation status ^24^, we proceeded to conduct Affymetrix analysis on Tel-treated ERα-positive BC cell lines to explore if additional kinases might be influenced by this drug and potentially involved in the regulation of receptor stability. For this purpose, we decided to undertake an unbiased approach by employing three different cell lines modelling the three major subtypes of ERα-positive breast tumors: MCF-7 cells were chosen because they represent the LumA phenotype, while BT-474 cells were selected because they belong the LumB class of BC. Finally, we also performed the same experiment in a cell line modelling a metastatic BC resistant to the ET drugs because they express an ERα missense mutation (i.e., Y537S) that renders the receptor hyperactive and sustains uncontrolled cell proliferation ^25,26^.

The results revealed that Tel administration significantly reduced the mRNA levels of 21 kinases in MCF-7 cells, 8 in BT-474 cells, and 44 in Y537S MCF-7 cells (FC ≤ -2; q-value ≤ 0.05) (Fig. 2A and Supplementary Table 3). Remarkably, only one kinase (CDK2) exhibited reduced levels in all three cell lines, while 17 kinases (BUB1, PLK1, DCLK1, CDC7, AURKB, CDK1, PBK, CAMK2D, CHK1, MELK, BUB1B, PLK4, GSG2, DYRK1B, VRK1, TTK, MASTL) were commonly reduced in both MCF-7 and Y537S MCF-7 cells. Intriguingly, we found that 12 (PLK1, CDC7, AURKB, CDK1, PBK, CHK1, MELK, BUB1B, PLK4, VRK1, TTK, MASTL) out of these 17 commonly reduced kinases (Fig. 2B and Supplementary Table 3) are part of a kinase signature that distinguishes LumA BC from basal BC ^27^. These findings suggest that Tel reduces the levels of several kinases in LumA BC cells, and this reduction may be linked to the degradation of ERα.

**Figure 2.**
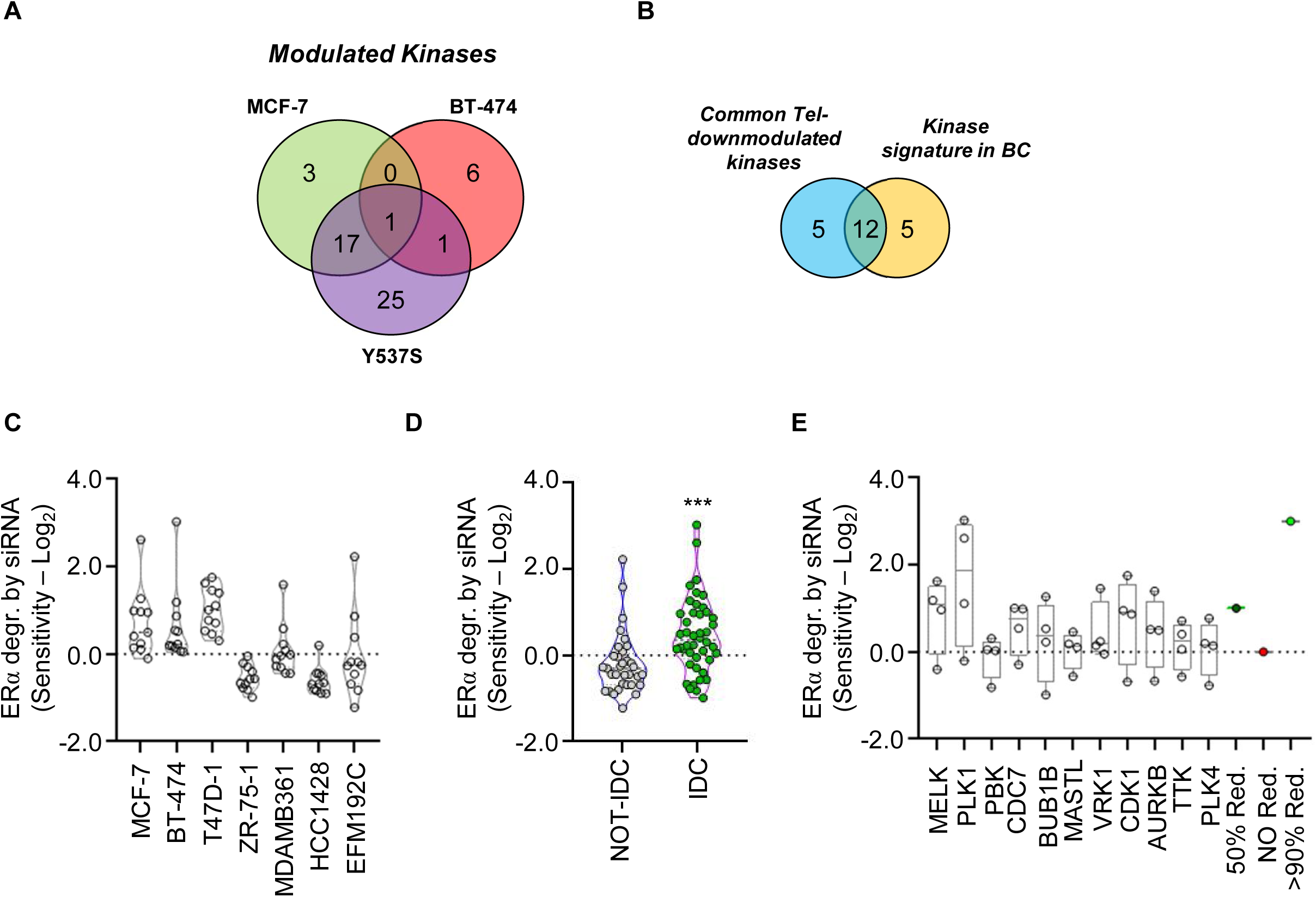
Potential Involvement of MELK Kinase in Regulating ERα Stability. (A) Venn diagram illustrating the number of modulated kinases (FC ≤ -2; q-value ≤ 0.05) as obtained through Affymetrix analyses in MCF-7, BT-474, and Y537S MCF-7 cells following a 24-hour administration of telaprevir (Tel -20µM). (B) Venn diagram displaying the kinases commonly modulated in MCF-7 and Y537S MCF-7 cells, along with the kinase signature identified in ^27^. (C) Sensitivity values in the indicated cell lines, reflecting the reduction in ERα intracellular levels assessed after treatment with esiRNA targeting the specific kinases identified in panel (B); each dot represents the value of a specific esiRNA. (D) Sensitivity values for reduction in ERα intracellular levels evaluated after treatment with esiRNA targeting the specific kinases identified in panel (B), stratified based on the histological type (invasive ductal carcinoma – IDC versus not-IDC) of the breast cancer (BC) cell lines used; each dot represents the value of a specific esiRNA. Statistical significance is indicated by *** (p<0.001) calculated using the Student-t test. (E) Sensitivity values for reduction in ERα intracellular levels assessed after treatment with esiRNA targeting the indicated kinases in IDC BC cell lines; each dot represents the value of the indicated esiRNA in the specific IDC cell line. For further details, please refer to the main text.

To test this hypothesis, we performed siRNA experiments using esiRNA reagents and we evaluated the impact of each esiRNA treatment on ERα content in the same abovementioned seven different cell lines. The experiments were repeated twice for screening purposes, and ERα levels were assessed 24 hours after the administration of esiRNA. To quantify the sensitivity of each treatment on ERα levels, we logarithmically transformed (-Log_2_) the fold of difference in ERα levels compared to controls for each esiRNA in each cell line. As shown in Fig. 2C and Supplementary Table 4, treatment with esiRNA targeting the 11 kinases (PLK1, CDC7, AURKB, CDK1, PBK, MELK, BUB1B, PLK4, VRK1, TTK, MASTL, excluding CHK1, as we had previously investigated its effect on the regulation of ERα levels and cell proliferation in different BC cell lines ^17^) was more effective in reducing ERα levels in MCF-7, BT-474, and T47D-1 cell lines than in ZR-75-1, MDA-MB-361, EFM192C, and HCC1428 cells. Notably, when we stratified the cell lines based on histological type (invasive ductal carcinoma – IDC *versus* not-IDC) ^18,19^, we observed that the reduction in ERα levels caused by esiRNA treatment against the 11 kinases was significantly overall higher in IDC cells (MCF-7, ZR-75-1, T47D-1, BT-474) than in not-IDC cells (HCC1428, EFM192C and MDA-MD-361) (Fig. 2D and Supplementary Table 4). Subsequently, we individually evaluated the effect of each esiRNA treatment in IDC cells and discovered that the depletion of PLK1 and MELK resulted in higher reductions in ERα levels (Fig. 2E). These findings suggest that treating IDC cell lines with esiRNA targeting several kinases, which are responsible for distinguishing the LumA BC phenotype from the basal BC phenotype ^27^, leads to a reduction in ERα levels. Furthermore, considering the known effect of PLK1 depletion on reducing ERα levels ^21,22^, we proceeded to conduct further investigations to examine the influence of MELK on the regulation of both ERα levels and cell proliferation.

### The impact of ALK and MELK in different BC subtypes

Subsequently, we investigated whether the effects of ALK and MELK on ERα stability were specific to particular subtypes of ERα-positive BC. For this purpose, we classified the aforementioned seven cell lines based on their PR and HER2 expression ^18,19^. Interestingly, the sensitivity for the reduction in ERα levels of the different cell lines to the esiRNA treatment against MELK was significantly higher in BC cell lines expressing PR (MCF-7, T47D-1, HCC1428, BT-474, and EFM192C) (Fig. 3A, 3C and Supplementary Table 4), while the sensitivity for the reduction in ERα levels of the different cell lines to AP26113 (AP)-dependent ALK inhibition was significantly higher in PR-negative cells (MDA-MB-361 and ZR-75-1 cells) (Fig. 3B, 3C and Supplementary Table 4).

**Figure 3.**
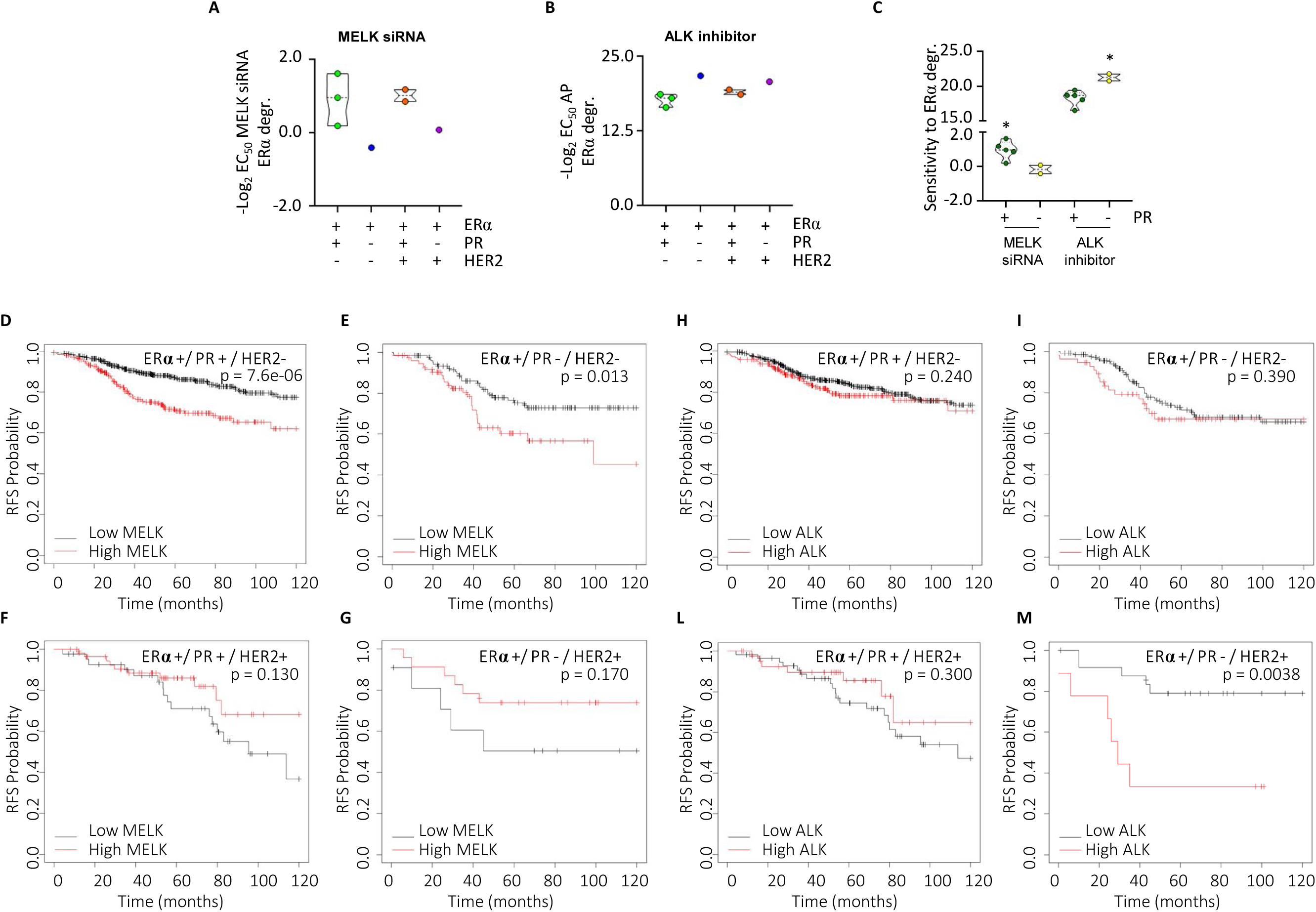
Breast Cancer Subtype Sensitivity to ALK and MELK Inhibition. (A) Sensitivity values in the indicated cell lines representing different breast cancer (BC) subtypes for reduction in ERα intracellular levels evaluated after treatment with esiRNA targeting MELK (A) or after administration of different doses of AP26113 – AP (B) and (C) stratified based on progesterone receptor (PR) expression. Statistical significance is indicated by * (p<0.05) calculated using the Student-t test. Kaplan-Meier plots showing the relapse-free survival (RFS) probability in women with breast tumors expressing different levels of ERα, progesterone receptor (PR), and HER2 in relation to MELK mRNA levels (D-G) or ALK mRNA levels (H-M). The p-values for significant differences between RFS are provided in each panel. Data obtained from the website (https://kmplot.com/analysis/). All possible cutoff values between the lower and upper quartiles are automatically computed (i.e., auto-select best cutoff on the website), and the best-performing threshold is used as a cutoff ^28^.

To further understand which BC phenotype could be more influenced by MELK and ALK expression, we examined the public KMplotter database (https://kmplot.com/analysis) ^28^ to assess the relapse-free survival (RFS) rate in women with ERα-positive BC, stratified based on PR and HER2 expression. The data revealed that women with low MELK mRNA levels displayed a significantly longer RFS rate than those with high MELK mRNA levels, particularly in tumors classified as ERα-positive/PR-positive/HER2-negative or ERα-positive/PR-negative/HER2-negative (Fig. 3D-3G and Supplementary Table 5), with the ERα-positive/PR-positive/HER2-negative phenotype showing the most significant difference. Conversely, women with low ALK mRNA levels displayed a significantly longer RFS rate than those with high ALK mRNA levels only in tumors classified as ERα-positive/PR-negative/HER2-positive (Fig. 3H-3M and Supplementary Table 5). These findings suggest that MELK could be a potential target in ERα-positive/PR-positive/HER2-negative BC cases, whereas ALK could be a target specifically in ERα-positive/PR-negative/HER2-positive tumors. Remarkably, these data align with the analysis conducted in the cell lines, supporting the notion that interference with MELK and ALK could affect ERα stability in BC cell lines stratified based on PR expression. Consequently, we selected MCF-7 and MDA-MB-361 cells, which display ERα-positive/PR-positive/HER2-negative and ERα-positive/PR-negative/HER2-positive phenotypes, respectively, to further validate the impact of these kinases on ERα stability and BC cell proliferation.

### Validation of the ALK and MELK impact on ERα levels and cell proliferation in MCF-7 and MDA-MB-361 cells

Subsequently, we validated the impact of esiRNA-mediated depletion and inhibition of both MELK and ALK on the intracellular content of ERα in MCF-7 and MDA-MB-361 cells. The results demonstrate that the depletion of MELK led to a substantial reduction in ERα levels solely in MCF-7 cells (Fig. 4A, 4A’, and 4A’’). Furthermore, treatment of MCF-7 and MDA-MB-361 cells with varying concentrations of the MELK inhibitor, MELK-8a (MELKin) ^29^, for 24 hours exhibited a dose-dependent decrease in ERα content in MCF-7 cells, whereas the effect on receptor levels in MDA-MB-361 cells was only marginal and observed at higher doses (10 µM) (Fig. 4B, 4B’, and 4B’’). esiRNA-mediated depletion of ALK in both cell lines resulted in a reduction of ERα content, which was significantly more pronounced in MDA-MB-361 cells compared to MCF-7 cells (Fig. 4C, 4C’, and 4C’’). Similarly, treatment of both cell lines with different doses of the ALK inhibitor AP, demonstrated a dose-dependent decrease in intracellular receptor content, with a more substantial effect observed in MDA-MB-361 cells (Fig. 4D, 4D’, and 4D’’).

**Figure 4.**
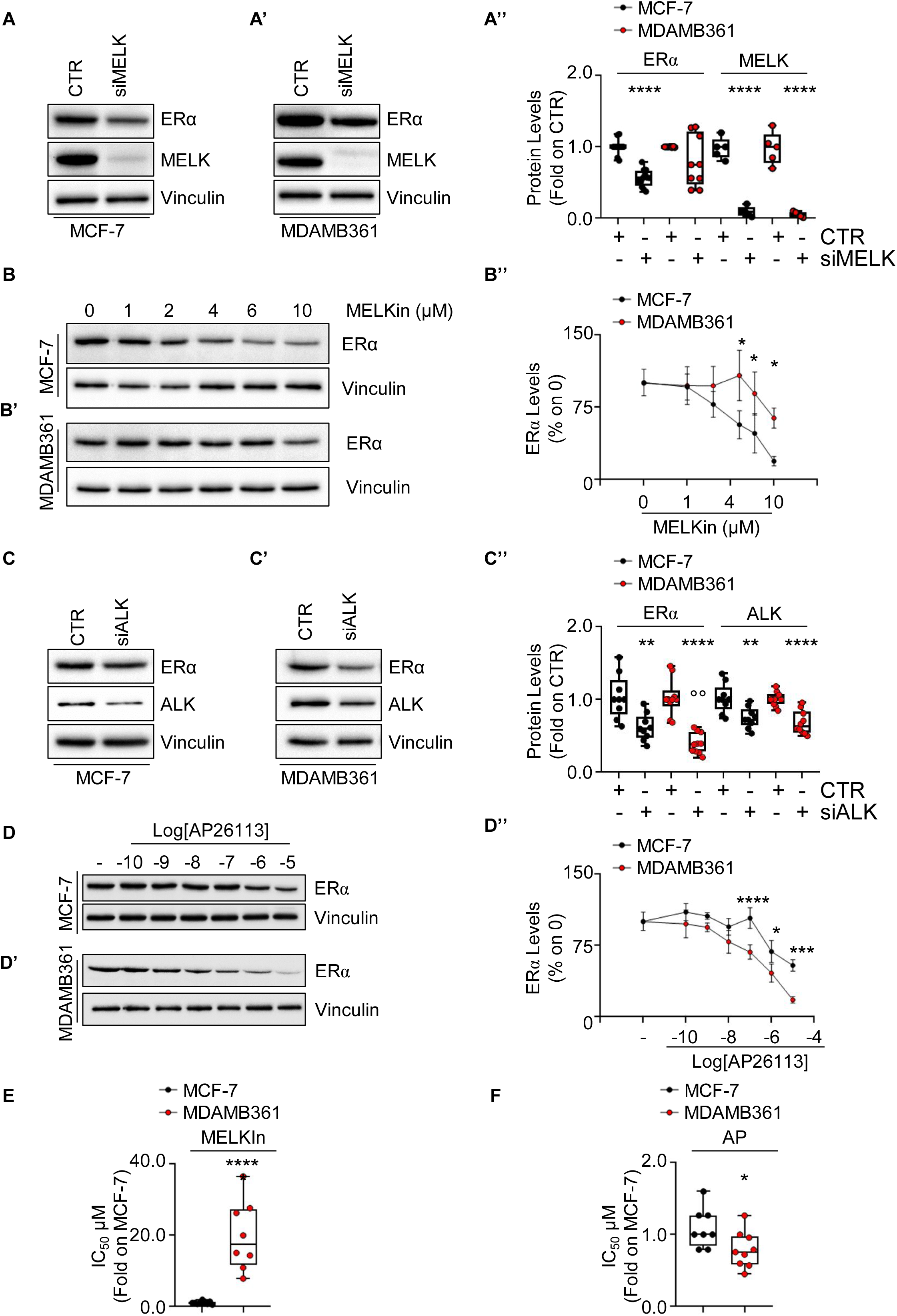
Confirmation of ALK and MELK Inhibition Effects on ERα Levels and Cell Proliferation in MCF-7 and MDA-MB-361 Cell Lines. Western blot analyses of ERα expression levels in MCF-7 and MDA-MB-361 cells treated with either MELK esiRNA (A, A’) or ALK esiRNA oligonucleotides for 24 hours (C, C’), as well as with indicated doses of the MELK inhibitor MELK-8a (MELKin) (B, B’) or the ALK inhibitor AP26113 (AP) (D, D’) for 48 hours. Representative blot images are shown. (A’’, B’’, C’’, and D’’) Densitometric analyses of the corresponding blots. In panels A’’ and C’’, significant differences were calculated using the ANOVA test, and * indicates differences compared to control (CTR) samples (**p<0.01, ****p<0.0001), while ° indicates differences compared to esiRNA-treated samples (°°p<0.01). In panels B’’ and D’’, significant differences were calculated for each dose in the different cell linesusing the Student-t test, and * represents a p-value < 0.05, *** represents p-values < 0.001, and **** represents p-values < 0.0001. (E) The inhibitor concentration 50 (IC_50_) calculated for both MCF-7 and MDA-MB-361 cells treated with different doses of the MELK inhibitor MELK8a (MELKin) for 7 days. Each dot represents an experimental replica. Significant differences were calculated using the Student-t test, and **** indicates a p-value < 0.0001. (F) The inhibitor concentration 50 (IC_50_) calculated for both MCF-7 and MDA-MB-361 cells treated with different doses of the ALK inhibitor AP26113 (AP) for 7 days. Each dot represents an experimental replica. Significant differences were calculated using the Student-t test, and * indicates a p-value < 0.05.

Subsequently, we evaluated the antiproliferative efficacy of MELKin and AP in both MCF-7 and MDA-MB-361 cells by generating growth curves and determining the inhibitory concentration 50 (IC_50_) for each compound in each cell line. Our findings revealed that the IC_50_ values for both cell lines fell within the µM range. Interestingly, the IC_50_ of MELKin in MCF-7 cells was significantly lower than that calculated in MDA-MB-361 cells. Conversely, the IC_50_ of AP in MDA-MB-361 cells was significantly lower than that observed in MCF-7 cells.

Collectively, these data indicate that interfering with MELK and ALK leads to a reduction in intracellular ERα content, thereby preventing BC cell proliferation. Furthermore, our results suggest that MELK predominantly controls ERα stability and cell proliferation in MCF-7 cells, while ALK more strongly modulates receptor intracellular levels and cell proliferation in MDA-MB-361 cells.

### The ALK- and MELK-dependent control of ERα intracellular concentration

Ligand-induced reduction of ERα in BC cells may result from the ligand’s ability to directly bind to the receptor ^15^. To examine this, ERα binding assays were conducted using various doses of AP, MELKin, and E2 to assess whether these kinase inhibitors could directly bind to ERα in vitro. Only E2 (Fig. 5A) was found to displace fluorescently labeled E2, used as a tracer for purified recombinant ERα, with an IC_50_ (i.e., K_d_) value of around 2.0 nM, consistent with previous reports ^10^. Next, the impact of kinase inhibition on ERα mRNA levels was investigated. Both MCF-7 and MDA-MB-361 cells were treated with MELKin and AP, respectively, for 48 hours. However, no significant difference in ERα mRNA content was observed in either cell line (Fig. 5B).

**Figure 5.**
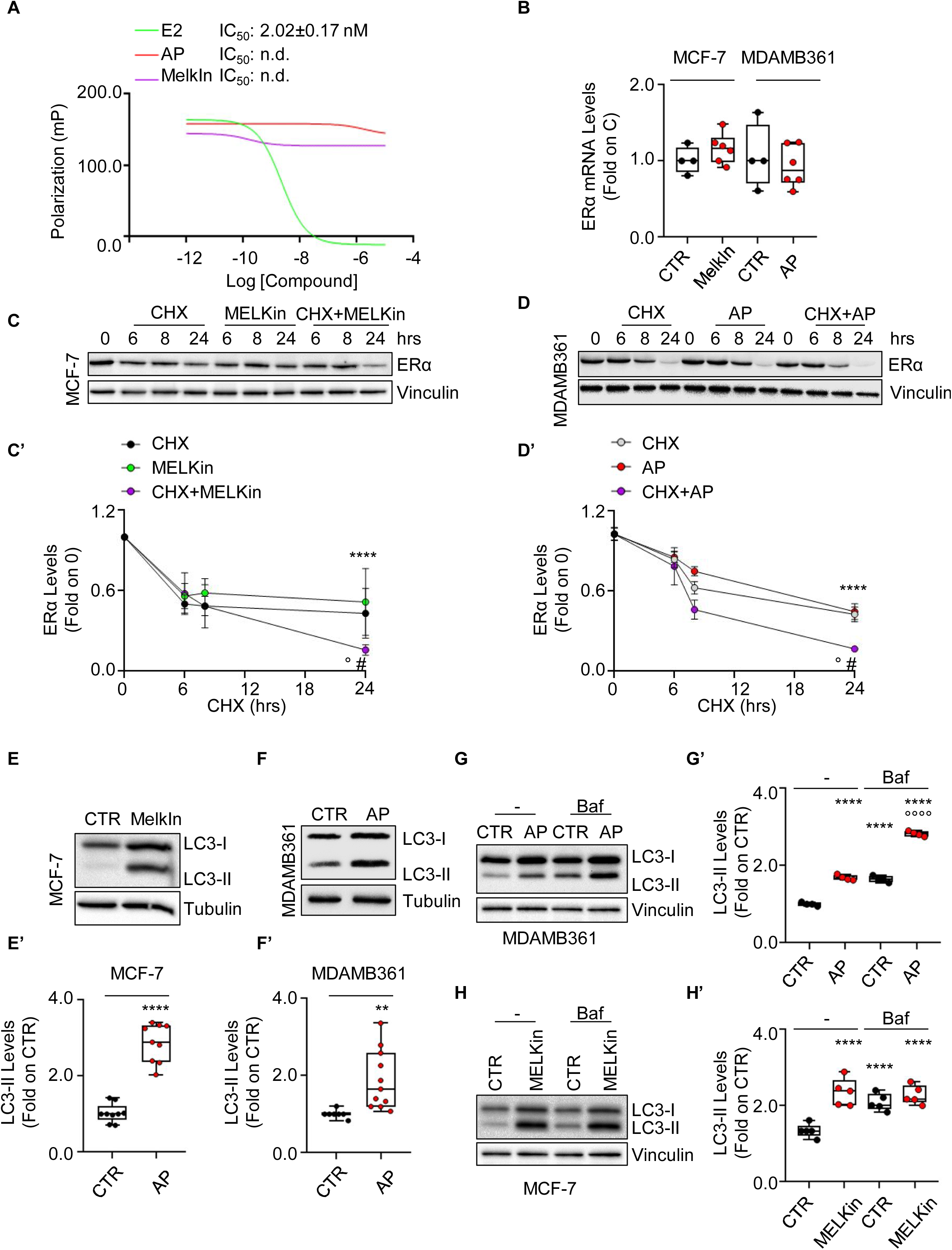
Mechanism of MELK and ALK Regulation on ERα Intracellular Levels in MCF-7 and MDA-MB-361 Cells. (A) In vitro ERα competitive binding assays were performed for the MELK inhibitor MELK-8a (MELKin), the ALK inhibitor AP26113 (AP), and 17β-estradiol (E2) at different compound doses, using fluorescent E2 as the tracer. The graph shows the relative inhibitor concentration 50 (IC_50_, i.e., K_d_) values. The experiment was conducted twice with five replicates. (B) Real-time qPCR analysis of ERα mRNA levels in MCF-7 cells treated for 24 hours with the MELK inhibitor MELK-8a (MELKin, and in MDA-MD-361 cells treated for 48 hours with the ALK inhibitor AP26113 (AP-1µM). The experiment was repeated twice with three replicates, and each dot represents an experimental replica. Western blot and relative densitometric analysis of ERα levels in MCF-7 cells (C) and in MDA-MB-361 cells (D) treated with cycloheximide (CHX - 1µM and 0.5µM, respectively) at different time points, both in the presence and absence of the MELK inhibitor MELK-8a (MELKin - 10µM) and the ALK inhibitor AP26113 (AP - 1µM). Representative blot images are shown. Significant differences with respect to the control (CTR) samples are calculated using the Student t-test and indicated by **** (p < 0.0001). Significant differences with respect to the CHX or inhibitor samples are calculated using the Student t-test and indicated by ° and # (p < 0.05), respectively. Western blot analysis and relative densitometric analyses of LC3 cellular levels in MCF-7 cells treated with the MELK inhibitor MELK-8a (MELKin - 10µM) (E, E’) and in MDA-MB-361 cells treated with the ALK inhibitor AP26113 (AP - 1µM) (F, F’) for 24 hours, both in the presence and absence of bafilomycin A1 (Baf - 100 nM) administration in the last 2 hours of treatment (G, G’, H, and H’). LC3 quantitation was performed using the formula LC3-II/(LC3-I+LC3-II). Representative blot images are shown. Significant differences with respect to the control (CTR) samples are calculated using the ANOVA test and indicated by **** (p < 0.0001). Significant differences with respect to the Baf samples are calculated using the ANOVA test and indicated by °°°° (p < 0.0001).

The turnover rate of ERα protein was then examined. MCF-7 and MDA-MB-361 cells were treated with the protein synthesis inhibitor cycloheximide (CHX) at different time points, both in the presence and absence of MELKin in MCF-7 cells and AP in MDA-MB-361 cells. As expected, MELKin, AP, and CHX reduced ERα levels. However, while CHX led to a time-dependent decay of the receptor, MELKin and AP effectively reduced ERα content only after 24 hours of treatment (Fig. 5C, 5C’, 5D, and 5D’). Interestingly, both inhibitors influenced the CHX-dependent reduction in ERα intracellular content after 24-hour administration (Fig. 5C, 5C’, 5D, and 5D’), suggesting that the kinase inhibitors can regulate ERα abundance at the post-translational level.

ERα stability can be modulated at the post-translational level through various cellular degradative pathways, such as the 26S proteasome, lysosomes, autophagic flux, and induction of replication stress ^8,17^. Therefore, we assessed the impact of each pathway on MELKin- and AP-induced reduction in ERα intracellular content both in MCF-7 and in MDA-MB-361 cells. We found that 24 hours administration of MELKin in MCF-7 cells and of AP in MDA-MB-361 cells determined the increase in the cellular amount of LC3-II [i.e., LC3-II/(LC3-I+LC3-II)], a marker of autophagosome number ^30^, thus indicating autophagosome accumulation (Fig. 5E, 5E’, 5F and 5F’). To determine whether this increase was due to autophagic flux activation or inhibition, additional experiments were conducted in the presence or absence of bafilomycin A1 (Baf), an inhibitor of the fusion between autophagosomes and lysosomes ^30^. In MDA-MB-361 cells, two hours of Baf administration resulted in increased LC3-II levels (Fig. 5G, 5G’), as expected ^30^. However, when Baf was added in the last two hours of AP treatment, it further significantly increased the levels of LC3-II compared to AP and Baf treatments alone (Fig. 5G, 5G’). Conversely, in MCF-7 cells, while two hours of Baf treatment increased LC3-II content (Fig. 5H, 5H’), adding Baf in the presence of MELKin did not further increase the amount of LC3-II levels induced by MELKin alone (Fig. 5H, 5H’). These findings indicate that AP activates autophagy in MDA-MB-361 cells, while MELKin inhibits the autophagic flux at its terminal stages in MCF-7 cells.

Taken together, these results indicate that ALK and MELK control ERα stability through a post-translational mechanism and regulate autophagy.

### The impact of MELK inhibition on E2:ERα signaling to cell proliferation

The ERα is a ligand-activated transcription factor that regulates the expression of multiple genes, both with and without the estrogen response element (ERE) sequence in their promoter regions in BC cells. Full E2-induced transcriptional activation of the receptor occurs upon phosphorylation of the S118 residue ^15^. Given the strong reduction in E2 signaling observed in cell lines modelling LumB BC ^31^, we investigated the impact of inhibiting MELK on E2 signaling and cell proliferation in MCF-7 cells. Upon E2 administration to MCF-7 cells, there was a notable increase in the phosphorylation of the S118 residue (Fig. 6A, 6A’, and 6A’’) as expected ^32^. Pretreatment of MCF-7 cells with MELKin or esiRNA-dependent depletion of MELK significantly reduced E2-induced ERα S118 phosphorylation (Fig. 6A, 6A’, and 6A’’). To study receptor transcriptional activity, we utilized MCF-7 cells stably expressing a reporter gene consisting of a promoter containing three synthetic ERE sequences that control the nanoluciferase gene (NLuc) (i.e., MCF-7NLuc cells) ^13^. E2 induced the activation of the synthetic ERE-containing promoter, and pretreatment with MELKin in MCF-7NLuc cells resulted in a dose-dependent reduction in E2-induced promoter activity (Fig. 6B). Moreover, depletion of MELK (inset in Fig. 6C) prevented the E2-dependent induction of ERE-containing promoter activity in MCF-7NLuc cells (Fig. 6C). As ERα controls the activation of genes with or without the ERE sequence in their promoter regions ^15^, we assessed the impact of MELK inhibition on E2-dependent gene expression. Using an RT-qPCR-based array containing 89 E2-sensitive genes ^7,23^, we hybridized cDNA samples generated from total RNA extracted from MCF-7 cells treated with E2 for 24 hours, both in the presence and absence of MELKin. As expected, most of the genes included in the array were modulated by E2 (i.e., 69.7%) (Fig. 6D). Interestingly, treatment with MELKin prevented the effect of E2 in 75.8% of the genes initially modulated by E2 in MCF-7 cells (Fig. 6D). Subsequently, we validated the effect of MELKin on some of these genes in MCF-7 cells. We pre-treated MCF-7 cells with MELKin and then treated them with E2, measuring the cellular levels of ERE-containing genes (presenilin 2 - pS2 and retinoic acid receptor A - RARA) and those lacking the ERE sequence in their promoter region (brain-derived nerve factor - BDNF and cyclin D1 - CycD1), along with the levels of ERα as an internal control. As expected, E2 induced an increase in the cellular levels of pS2, RARA, BDNF, and CycD1 and led to ERα degradation after 24 hours of administration to MCF-7 cells (Fig. 6E-6M). Notably, inhibition of MELK, as well as reduction in MELK expression, prevented the E2-induced increase in pS2, RARA, BDNF, and CycD1 expression levels and resulted in an additional reduction in the receptor’s intracellular content (Fig. 6E-6M). Collectively, these data indicate that MELK inhibition decreases ERα transcriptional activity, impedes E2’s ability to activate ERα, and hinders E2-dependent gene expression.

**Figure 6.**
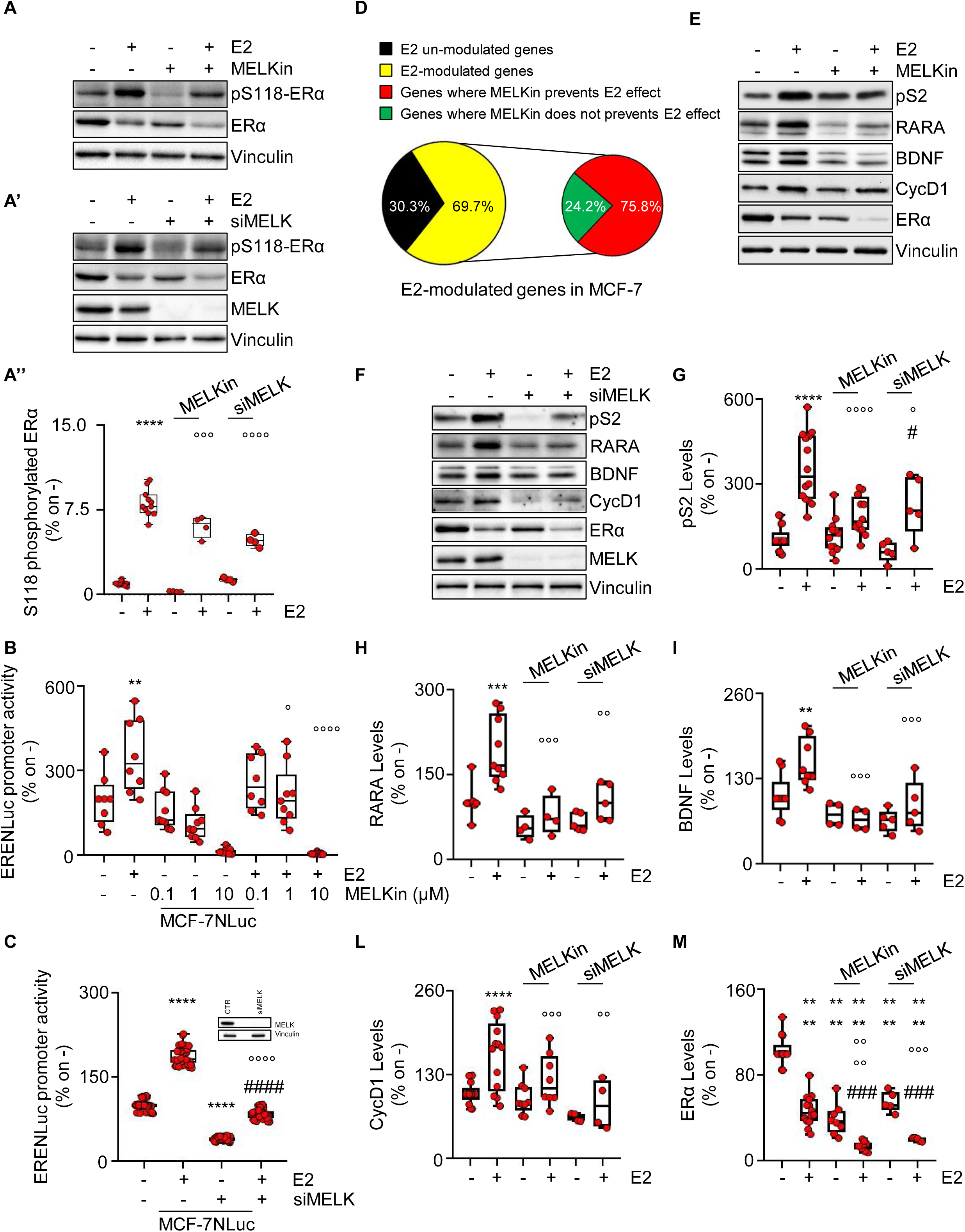
MELK Inhibition Impacts E2:ERα Transcription Signaling in MCF-7 Cells. (A) Western blot and relative densitometric analyses of ERα and ERα S118 phosphorylation expression levels in MCF-7 cells pre-treated with the MELK inhibitor MELK-8a (MELKin - 10µM) for 24 hours (A, A’’) or with MELK esiRNA (A’ and A’’) and then treated for 30 minutes with 17β- estradiol (E2 - 1 nM). Representative blot images are shown. Significant differences with respect to the untreated (-) sample are calculated using the ANOVA test and indicated by **** (p-value < 0.0001). Significant differences with respect to the E2-treated sample are calculated using the ANOVA test and indicated by °°° (p-value < 0.001) or °°°° (p-value < 0.0001). (B) Estrogen response element promoter activity in MCF-7 ERE-NLuc cells pre-treated with the MELK inhibitor MELK-8a (MELKin - 10µM) for 24 hours (B) or with MELK esiRNA (C) and then treated with 17β-estradiol (E2 - 1 nM) for an additional 24 hours. The experiments were performed three times in quintuplicate. Significant differences were calculated using the ANOVA test. ** (p-value < 0.01) and **** (p-value < 0.0001) indicate significant differences with respect to the untreated (-) sample. ° (p-value < 0.05) and °°°° (p-value < 0.0001) indicate significant differences with respect to the E2-treated sample. #### (p-value < 0.0001) indicates significant differences with respect to the MELK esiRNA-treated sample. (C) Pie diagrams illustrating the percentages of modulated array genes in MCF-7 cells pre-treated with the MELK inhibitor MELK-8a (MELKin - 10µM) for 24 hours and then treated with 17β-estradiol (E2 - 1 nM) for an additional 24 hours. Percentages and categories of genes are indicated. (D) Western blot of presenilin 2 (pS2), retinoic acid receptor A (RARA), brain-derived nerve factor (BDNF), cyclin D1 (CycD1), and ERα expression levels in MCF-7 cells pre-treated with the MELK inhibitor MELK-8a (MELKin - 10µM) for 24 hours (E) or with MELK esiRNA (F) and then treated with 17β-estradiol (E2 - 1 nM) for an additional 24 hours. Representative blot images are shown. Densitometric and statistical analyses are reported for each protein in panels F-M. Significant differences were calculated using the ANOVA test. **, *** and **** indicate significant differences with respect to the untreated (-) sample. °, °°, °°° and °°°° (p-value < 0.05, < 0.01, 0.001 and < 0.0001, respectively) indicate significant differences with respect to the E2-treated sample. ### (p-value < 0.001) indicates significant differences with respect to the MELKin or MELK esiRNA-treated samples. Each dot represent an experimental replica.

Since E2-dependent activation of ERα in BC cells leads to DNA synthesis, cell cycle progression, and cell proliferation ^15^, we investigated the effect of MELK inhibition on E2’s ability to induce these processes in MCF-7 cells. Treatment with both MELKin and esiRNA targeting MELK (inset in Fig. 7A) significantly reduced E2-induced 5-ethynyl-2’-deoxyuridine (EdU) incorporation in MCF-7 cells (Fig. 7A). Furthermore, E2 increased the cell number in a time-dependent manner, and co-treatment of MCF-7 cells with MELKin prevented both the basal and E2-induced time-dependent increase in cell number (Fig. 7B).

**Figure 7.**
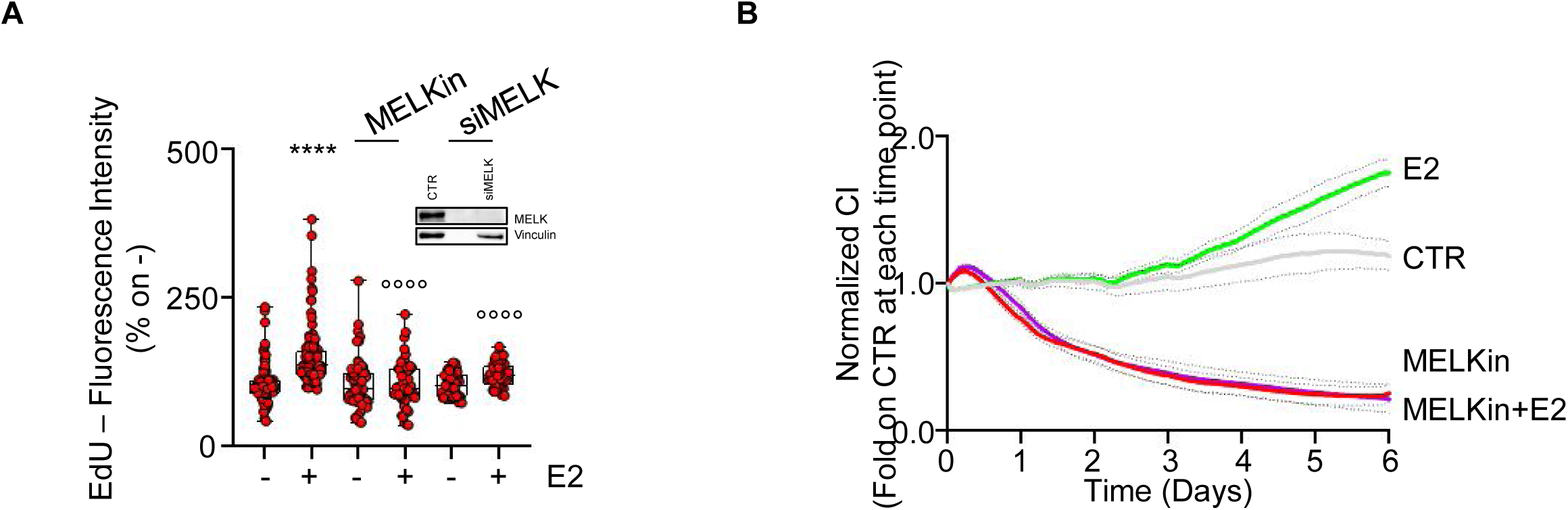
Impact of MELK Inhibition on E2-Induced Cell Proliferation in MCF-7 Cells. (A) 5-ethynyl-2’-deoxyuridine (EdU) incorporation assay in MCF-7 cells treated with 17β-estradiol (E2 - 1 nM) for 24 hours after 24 hours pre-treatment with the MELK inhibitor MELK-8a (MELKin - 10µM) or with MELK esiRNA. The experiments were performed twice in quintuplicate. Significant differences were calculated using the ANOVA test. **** (p-value < 0.0001) indicates significant differences with respect to the untreated (-) sample. °°°° (p-value < 0.0001) indicates significant differences with respect to the E2-treated sample. (B) The graphs show the normalized cell index (i.e., cell number) detected with the xCelligence DP device and calculated at each time point with respect to the control sample. Each sample was measured in quadruplicate. MCF-7 cells were treated with 17β-estradiol (E2 - 1 nM) and the MELK inhibitor MELK-8a (MELKin - 10µM) when cells were plated. Dotted lines represent standard deviations.

Altogether, these findings indicate that inhibition of MELK activity interferes with E2’s ability to induce DNA synthesis and cell proliferation in MCF-7 cells.

### MELK and ALK inhibitors in combination with 4OH-tamoxifen and HER2 inhibitors as a novel selective treatment for specific BC subtypes

The obtained results suggest that MELK could serve as a promising target for treating ERα- positive breast tumors with the ERα-positive/PR-positive/HER2-negative phenotype. Conversely, our findings indicate that ALK could be targeted in tumors with the ERα-positive/PR-negative/HER2-positive phenotype. It is worth noting that tumors with the ERα-positive/PR-positive/HER2-negative phenotype are typically treated with Tam, the standard treatment for this type of tumors ^4,5^, while HER2-positive tumors are treated with drugs inhibiting HER2 activity (e.g., lapatinib – Lapa, erlotinib – Erlo, and gefitinib – Gef) ^4,5^. Therefore, we sought to investigate whether combining MELKin with Tam and combining the ALK inhibitor AP with Lapa, Erlo, and Gef could have potential benefits in MDA-MB-361 cells. Proliferation studies were performed by treating cells for 12 days with varying doses of MELKin together with varying doses of Tam in MCF-7 cells and different doses of Lapa, Erlo, and Gef along with different doses of AP in MDA-MB-361 cells. The data reveal that Tam and MELKin synergistically enhance the anti-proliferative effects of both inhibitors in MCF-7 cells (Fig. 8A, 8A’). Interestingly, while AP synergistically enhances the effect of all HER2 inhibitors in MDA-MB-361 cells (Fig. 8B-E), we observed that the combination of AP with either Erlo or Gef was more effective than the combination of AP and Lapa in achieving an anti-proliferative effect in MDA-MB-361 cells (Fig. 8B-E).

**Figure 8.**
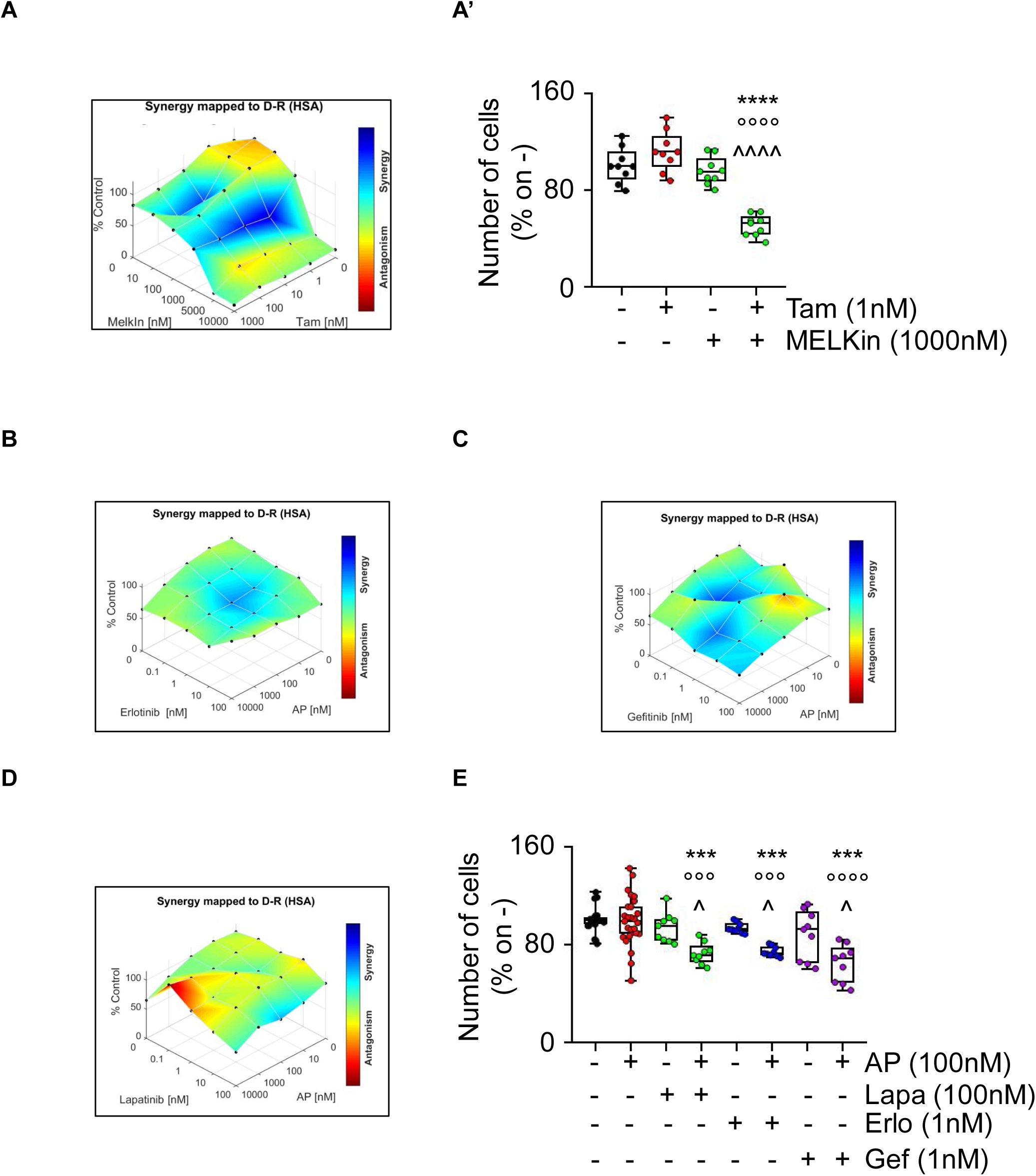
Synergy between MELK and 4OH-Tamoxifen in MCF-7, and between ALK and HER2 Inhibitors in MDA-MB-361 Cells. (A) Synergy map of 12-day-treated MCF-7 cells with different doses of 4OH-Tamoxifen (Tam) and the MELK inhibitor MELK-8a (MELKin). (B’) Growth curves in MCF-7 cells showing the synergistic effect of each combination of compounds with selected doses. Significant differences were calculated using the ANOVA test. **** (p-value < 0.0001) indicates significant differences with respect to the untreated (i.e., -,-) sample. °°°° (p-value < 0.0001) indicates significant differences with respect to Tam treated sample. ^^^^ (p-value < 0.0001) indicates significant differences with respect to MELKin treated sample. Synergy map of 12-day-treated MDA-MB-361 cells with different doses of the ALK inhibitor AP26113 (AP) and the HER2 inhibitors erlotinib (Erlo) (B), gefitinib (Gef) (C), and lapatinib (Lapa) (D). (E) Growth curves in MDA-MB-361 cells showing the synergistic effect of each combination of compounds with selected doses. Significant differences were calculated using the ANOVA test. *** (p-value < 0.001) indicates significant differences with respect to the untreated (i.e., -,-) sample. °°°, °°°° (p-value < 0.001 and < 0.0001, respectively) indicates significant differences with respect to Erlo, Gef, and Lapa treated samples. ^ (p-value < 0.05) indicates significant differences with respect to AP treated sample.

These findings support the concept that MELKin could be a promising candidate for combinatorial treatment in ERα-positive/PR-positive/HER2-negative tumors in conjunction with Tam, and the ALK inhibitor AP could be considered for combinatorial treatment in ERα-positive/PR-negative/HER2-positive tumors with HER2 inhibitors.

### Evaluation of the antiproliferative effect of MELK and ALK inhibitors in 3D models of BC

We finally investigated the anti-proliferative effects of MELKin and AP in MCF-7 and MDA-MB-361 tumor cell spheroids and alginate-based cultures ^16,17^ to assess their activity in 3D cell structures ^33^. Both tumor spheroids and cells within alginate-based spheres demonstrated successful growth within a 7-day period. Remarkably, treatment with MELKin significantly inhibited cell proliferation in both MCF-7 spheroids and alginate-based structures (Fig. 9A, 9A’, 9B, and 9B’). However, in the case of MDA-MB-361 cells, while AP administration effectively prevented proliferation in alginate-based spheres, it had no significant effect on cell growth when the cells were cultured as spheroids (Fig. 9A, 9A’, 9B, and 9B’). These results indicate that MELKin and AP retain their anti-proliferative efficacy in 3D models of BC, although they may exert their effects through distinct mechanisms of action.

**Figure 9.**
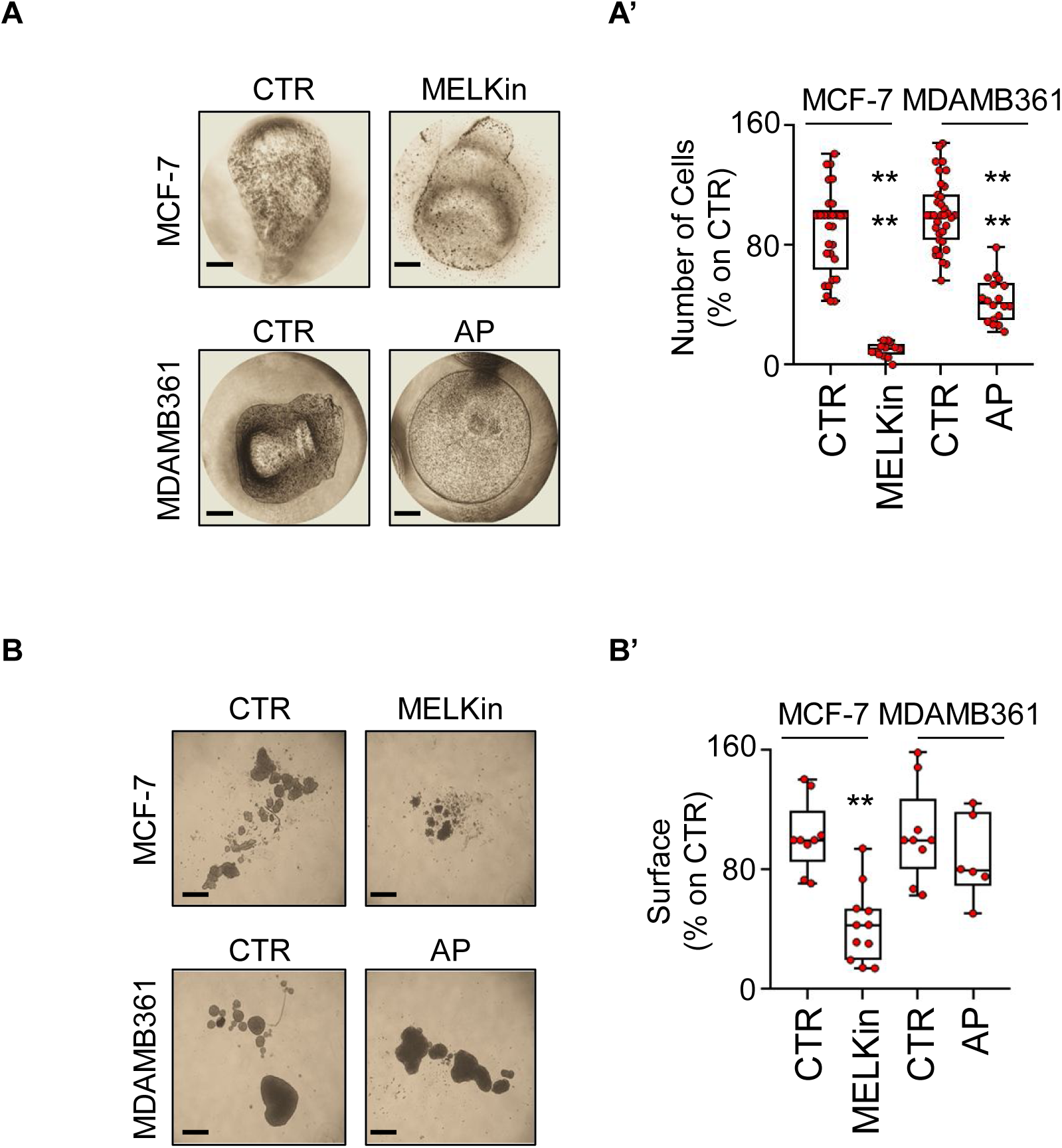
Effect of MELK and ALK Inhibitors in 3D Models of Breast Cancer. Images (A, B) and quantitation (A’, B’) of tumor spheroids’ surface area (B, B’) and alginate-based cultures (A, A’) generated in MCF-7 and MDA-MB-361 cells, treated at time 0 with the MELK inhibitor MELK-8a (MELKin - 10µM), the ALK inhibitor AP26113 (AP - 1µM), or left untreated (CTR) for 7 days. The number of replicates is represented by solid dots in the graphs. Significant differences with respect to the CTR sample were determined using unpaired two-tailed ANOVA test: **** (p-value < 0.0001); ** (p-value < 0.01). Scale bars equal to 50.0 µm.

## Discussion

The classification of breast cancer (BC) at diagnosis plays a critical role in determining the pharmacological approach for treating the disease. BC classification is based on various molecular and histological prognostic factors. The expression of ERα categorizes the tumor into two groups, each of which can be further stratified based on the histological type of the disease and the expression of PR and HER2. Additionally, PAM50 analysis of breast tumors identifies the luminal (LumA and LumB) or basal origin of the disease ^1–4^. Notably, specific breast tumor types can be a combination of all these factors, resulting in a unique tumor type for each patient, which may even be considered a rare disease ^34^. The heterogeneity of BC necessitates specific drugs that can selectively target particular BC subtypes to implement a personalized medicine approach. Notably, endocrine therapy drugs like Tam exhibit increased sensitivity in LumA tumors compared to LumB tumors, as the latter express HER2, which is better targeted by its inhibitors (erlotinib, gefitinib, lapatinib) ^4,5^. Consequently, identifying drugs that can selectively target specific tumor subtypes becomes increasingly important for effective BC treatment.

Recently, we found that certain drugs not originally intended for this purpose can induce receptor degradation in ERα-positive BC cells, making them act as ‘anti-estrogen-like’ compounds to prevent cell proliferation ^6–14^. Additionally, some of these drugs selectively induced ERα degradation and prevented cell proliferation only in specific BC subtypes ^6–14^. This led us to hypothesize that ERα-positive BC cells may be more sensitive to certain drugs than ERα-negative BC cells due to these compounds’ ability to induce ERα degradation, thereby displaying a selective effect on specific BC subtypes.

Taking advantage of sensitivity data from over 4600 drugs tested against 26 different ERα-positive and ERα-negative BC cell lines available in the DepMap portal database (https://depmap.org/portal/), we identified a list of 73 drugs that exhibited increased sensitivity in ERα-positive BC cell lines compared to ERα-negative ones. Among these drugs, we discovered 2 anti-helminthics compounds, 4 cardiac glycosides (CG), and 7 DNA polymerase inhibitors, known to produce replication stress ^20^. Interestingly, our recent findings also demonstrated that anti-helminthics clotrimazole and fenticonazole bound to ERα, induced its degradation, and prevented the proliferation of ERα-positive BC cell lines ^7^. Moreover, we reported that the CG compounds ouabain and digoxin showed increased sensitivity in ERα-positive BC cancer cell lines compared to ERα-negative BC cell lines because, in addition to inhibiting the Na/K ATPase, they hyperactivated the 26S proteasome, inducing receptor degradation ^6,10^. Additionally, we showed that CHK1 inhibitors induced replication stress, leading to ERα degradation ^17^. Therefore, the identified list of drugs could contain molecules capable of inducing ERα degradation.

Remarkably, 37 out of the 73 drugs on the list are kinase inhibitors, prompting us to focus on this class of molecules as kinases represent excellent drug targets controlling various pathways required for cell proliferation ^35^. Most of the identified kinase inhibitors targeted CHK1 and PLK1, which have been previously shown to induce receptor degradation ^17,21,22^. We also found 4 inhibitors in the list for AURKA/AURKB and ALK, and their impact on BC cell proliferation was poorly investigated. To address this, we studied whether the inhibition of these kinases could influence the cellular amount of ERα in 7 different ERα-positive BC cell lines representing different BC molecular and histological subtypes ^18,19^. We observed that ALK inhibitors led to a reduction in ERα levels.

We also used a hypothesis-driven approach to identify additional kinases involved in regulating receptor intracellular levels by conducting Affymetrix analyses on ERα-positive BC cells treated with telaprevir (Tel), an antiviral drug inducing ERα degradation by inhibiting the kinases IGF1-R and AKT ^23,24^. Surprisingly, we found that Tel reduced the mRNA levels of many kinases, most of which belonged to the kinase signature that distinguishes LumA BC from basal BC ^27^. We then tested the impact of reducing each of these kinases on ERα levels in the aforementioned BC cell lines and found that the reduction of receptor levels caused by cell treatment with esiRNA directed against these kinases was predominant in invasive ductal carcinoma (IDC) cells compared to not-IDC cells. Remarkably, we also observed that, in addition to PLK1, only the treatment with esiRNA directed against MELK led to a reduction in ERα levels.

Due to the lack of information on ALK- and MELK-dependent control of ERα levels, we further studied the impact of these two kinases in BC. We stratified sensitivity data for the reduction in ERα levels based on the expression of PR and HER2 in ERα-positive cell lines used and observed that cell lines expressing PR were more sensitive to the reduction in ERα levels induced by esiRNA directed against MELK, while cells not expressing PR were more susceptible to the ALK inhibitor AP26113 (AP)-dependent reduction in receptor levels. Accordingly, we found that low MELK and ALK mRNA expression is associated with a significantly improved patient RFS rate, depending on whether the patient carries a tumor with the ERα-positive/PR-positive/HER2-negative or the ERα-positive/PR-negative/HER2-positive phenotype, respectively. Thus, to investigate ALK and MELK’s impact, we studied MELK and ALK in ERα-positive BC cell lines showing the corresponding phenotype (i.e., MCF-7 and MDA-MB-361 cells, respectively ^18,19^). Using these cell lines, we demonstrated that MELK inhibition or depletion preferentially affected the control of ERα levels and cell proliferation in LumA, IDC, PR-positive and HER2-negative MCF-7 cells. In this cell line, interference with MELK activity or levels also prevented the receptor’s ability to control E2-induced transcriptional activity, gene expression, DNA synthesis, and cell proliferation. Conversely, ALK inhibition or depletion selectively affected the control of ERα levels and cell proliferation in LumB, adenocarcinoma, PR-negative, and HER2-positive MDA-MB-361 cells. However, we could not measure the ERα signaling to cell proliferation in this cell line, as E2 has a negligible effect on LumB cell lines ^31^.

Regarding the mechanism through which ERα is degraded upon ALK and MELK inhibition, we found that it occurs at post-translational level and does not imply the ability of the ALK and MELK inhibitors either to directly bind to the receptor or to control the ERα mRNA levels. However, we found that treatment with the MELK inhibitor blocked autophagy in MCF-7 cells, while the ALK inhibitor AP induced autophagy in MDA-MB-361 cells.

Previous data from our lab demonstrated that autophagic flux controls basal ERα degradation, and ERα is partially degraded in autophagosomes. Therefore, the effect induced by ALK and MELK inhibitors on the regulation of receptor intracellular levels could occur at post-translational levels through the modulation of the autophagic flux. Accordingly, in MDA-MB-361 cells, ALK inhibitor AP administration induced autophagy and resulted in receptor degradation. Surprisingly, in MCF-7 cells, the MELK inhibitor-induced ERα degradation was accompanied by autophagic flux inhibition. Two possibilities exist to explain this contradiction. ERα binds to p62^SQSTM^ and is shuttled to the autophagosomes by p62^SQSTM^ ^36^. Interestingly, p62^SQSTM^ plays a critical role in the balance between autophagic flux and the ubiquitin-proteasome system (UPS). Autophagy inhibition with increased p62^SQSTM^ levels has been reported to deregulate p62^SQSTM^-dependent shuttling of ubiquitinated proteins to the 26S proteasome ^37,38^. Therefore, it is tempting to speculate that in MCF-7 cells treated with the MELK inhibitor, ERα is degraded through the UPS via increased p62^SQSTM^-dependent shuttling to the proteasome. Additionally, in MCF-7 cells, a similar situation occurs under E2 administration, as E2 blocks autophagic flux and induces ERα degradation ^36^. The steady-state cellular ERα content is influenced by degradative pathways acting on both neo-synthesized and mature ERα fractions ^39^. We have shown that E2 impedes autophagic degradation of neo-synthesized ERα without affecting autophagy’s impact on the mature receptor pool ^36^. Therefore, it is also possible that MELK inhibitor-induced autophagy inhibition differentially affects the neo-synthesized and mature ERα pools. However, our data suggest that the autophagic control of ERα levels can follow different routes in different cell lines and this differential mechanistic aspect is currently being evaluated. Furthermore, our results indicate that both ALK and MELK are involved in controlling autophagy. Altogether, this evidence demonstrates that MELK and ALK control ERα stability and cell proliferation selectively in different BC subtypes.

Due to the differential effects observed in cell lines modeling various BC subtypes, we further evaluated the potential use of MELK and ALK inhibitors in pre-clinical combinatorial studies with drugs used to treat specific patient tumor phenotypes, including ERα-positive/PR-positive/HER2-negative and ERα-positive/PR-negative/HER2-positive phenotypes. Although the MELK inhibitor MELK-8a is not approved for use in humans, we observed that this drug exhibited a synergic antiproliferative effect when used in combination with Tam in MCF-7 cells. On the other hand, the ALK inhibitor AP, in clinical trials for patients with lung tumors ^40^, showed synergy with HER2 inhibitors, with varying effectiveness when co-administered with gefitinib and erlotinib compared to lapatinib. These results demonstrate that MELK inhibition could be a valuable strategy for treating BC patients with the ERα-positive/PR-positive/HER2-negative phenotype, while ALK inhibition, in combination with specific HER2 inhibitors, could be effective for treating ERα-positive/PR-negative/HER2-positive BC patients. Finally, the use of MELK and ALK inhibitors in BC patients is further supported by the fact that these inhibitors retained their anti-proliferative activities, albeit with some differences, in 3D models of BC, which mimic a context closer to the tumor environment ^33^.

## Conclusions

In this study, we present new findings identifying MELK and ALK as promising targets for the treatment of ERα-positive BC. Notably, we have uncovered that distinct BC subtypes, namely ERα-positive/PR-positive/HER2-negative and ERα-positive/PR-negative/HER2-positive, exhibit selective sensitivity to the inhibition of these kinases, respectively. Our research further demonstrates that targeting ERα-positive cells with the ERα-positive/PR-positive/HER2-negative receptor profile using the MELK inhibitor alone or in combination with the endocrine therapy drug Tam, as well as targeting ERα-positive cells representing the ERα-positive/PR-negative/HER2-positive phenotype with the ALK inhibitor AP alone or in combination with HER2 activity-blocking drugs such as gefitinib and erlotinib, offer promising strategies to curb the cell proliferation of specific ERα-positive BC subtypes.

Consequently, we propose that the targeted inhibition of MELK and ALK using small molecules could hold significant potential for personalized BC management. These findings may pave the way for more effective and tailored treatments for individuals with ERα-positive BC, offering new avenues for precision medicine in this context.

## Methods

### Cell Culture and Reagents

The following cell lines and chemicals were used: MCF-7, T47D-1, ZR-75-1, HCC1428, BT-474, and MDA-MB-361 cell lines were obtained from ATCC (USA), while EFM192C cells were obtained from DSMZ (Braunschweig, Germany). All cell lines were maintained according to the manufacturer’s instructions. The following reagents and antibodies were used: 17β-estradiol (E2), DMEM (with and without phenol red), and fetal calf serum were purchased from Sigma-Aldrich (St. Louis, MO). The Bradford protein assay kit, anti-mouse, and anti-rabbit secondary antibodies were obtained from Bio-Rad (Hercules, CA). Antibodies against ERα (F-10, mouse), pS2 (FL-84, rabbit), cyclin D1 (H-295 rabbit), ALK (F-12, mouse), and RARA (C-1, mouse) were obtained from Santa Cruz Biotechnology (Santa Cruz, CA, USA). Additionally, anti-MELK (ab273015, rabbit) and anti-BDNF (ab108319, rabbit) antibodies were purchased from Abcam (Cambridge, UK). Anti-phospho ERα (Ser118, mouse) antibody was obtained from Cell Signaling, and anti-vinculin (mouse) and anti-LC3 (mouse) antibodies were purchased from Sigma-Aldrich (St. Louis, MO, USA). Chemiluminescence reagent for Western blot was obtained from BioRad Laboratories (Hercules, CA, USA). For specific experiments, the following compounds were used: 4OH-Tamoxifen, cycloheximide (CHX), and esiRNA library were purchased from Sigma-Aldrich (St. Louis, MO, USA). MELK-8a hydrochloride, TAK-901, AT-9283, CCT-137690, AP26113, NVP-TAE-684, AZD-3463, Lapatinib, Gefitinib, and Erlotinib were purchased from Selleck Chemicals (USA). The PolarScreen™ ERα Competitor Assay Kit, Green (A15882) was acquired from Thermo Scientific. All other products used were from Sigma-Aldrich, and analytical- or reagent-grade products were used without further purification. To verify the authenticity of the cell lines, STR analysis was performed by BMR Genomics (Italy).

### In Vitro ERα Binding Assay

The in vitro ERα binding assay employed a fluorescence polarization (FP) method to assess the binding affinity of MELK-8a hydrochloride, AP26113, and 17β-estradiol (E2) with recombinant ERα. The FP assay was conducted using the PolarScreen™ ERα Competitor Assay Kit, Green (A15882, Thermo Scientific), following established procedures described in ^17^.

### Measurement of ERα Transcriptional Activity

The ERα transcriptional activity was assessed by measuring the expression of nanoluciferase (NLuc)-PEST, a reporter gene containing an estrogen response element (ERE), in stably transfected MCF-7 cells. After 24 hours of compound administration, the NLuc-PEST expression was determined following the described procedure ^13,41^.

### Cell Manipulation for Western Blot Analyses

Cells were initially cultured in DMEM containing phenol red and 10% fetal calf serum for 24 hours. Subsequently, the cells were treated with various compounds at specified doses and time periods as indicated. Before E2 stimulation, cells were cultured in DMEM without phenol red and 10% charcoal-stripped fetal calf serum for 24 hours. Addition of MELK8a occurred 24 hours prior to E2 administration. Following the treatments, cells were lysed in Yoss Yarden (YY) buffer, which consisted of 50 mM Hepes (pH 7.5), 10% glycerol, 150 mM NaCl, 1% Triton X-100, 1 mM EDTA, and 1 mM EGTA, supplemented with protease and phosphatase inhibitors. For Western blot analysis, 20–30 µg of protein was loaded onto SDS-gels. Gels were run, and the proteins were transferred to nitrocellulose membranes using a Turbo-Blot semidry transfer apparatus from Bio-Rad (Hercules, CA, USA). Immunoblotting was performed by incubating the membranes with 5% milk or bovine serum albumin for 60 minutes, followed by overnight incubation with the designated antibodies. Subsequently, secondary antibody incubation was carried out for an additional 60 minutes. Finally, the protein bands were detected using a Chemidoc apparatus from Bio-Rad (Hercules, CA, USA).

### Small Interference RNA

For the small interference RNA (siRNA) experiments, cells were transfected with esiRNA targeting the specific proteins of interest. The transfection procedure was conducted using Lipofectamine RNAi Max (Thermo Fisher), following established protocols described in ^42^.

### Cell Proliferation and 3D Cell Culture Assays

The xCELLigence DP system (ACEA Biosciences, Inc., San Diego, CA) Multi-E-Plate station was utilized to measure the time-dependent response to the specified drugs by real-time cell analysis (RTCA), following previously reported protocols ^10,13,17,23^. Synergy studies were conducted using Crystal Violet staining, as described in ^43^. The synergy was subsequently calculated using Combenefit freeware software ^17^. Alginate-based and tumor spheroid cultures were carried out following established procedures as previously reported ^17^.

### RNA isolation and qPCR analysis

Gene-specific forward and reverse primers were designed using the OligoPerfect Designer software program (Invitrogen, Carlsbad, CA, USA). For human ERα, the primers used were 5’-GTGCCTGGCTAGAGATCCTG-3’ (forward) and 5’-AGAGACTTCAGGGTGCTGGA-3’

(reverse). For human GAPDH, the primers used were 5’-CGAGATCCCTCCAAAATCAA-3’ (forward) and 5’-TGTGGTCATGAGTCCTTCCA-3’ (reverse). Total RNA was extracted from the cells using TRIzol Reagent (Invitrogen, Carlsbad, CA, USA), following the manufacturer’s instructions. For gene expression analysis, cDNA synthesis and qPCR were performed using the GoTaq 2-step RT-qPCR system (Promega, Madison, MA, USA) with an ABI Prism 7900HT Sequence Detection System (Applied Biosystems, Foster City, CA, USA), according to the manufacturer’s instructions. Each sample was tested in triplicates, and the experiment was repeated twice to ensure accuracy and reproducibility. Gene expression levels were normalized to GAPDH mRNA levels as an internal control.

### Gene Arrays Analyses

Gene Arrays Analyses were conducted as follows: Total RNA was extracted from cells using TRIzol reagent (Invitrogen, Carlsbad, CA, USA) following the manufacturer’s guidelines. For gene expression analysis, the GoTaq 2-step RT-qPCR system (Promega, Madison, MA, USA) was utilized to perform cDNA synthesis and qPCR. The ABI Prism 7900 HT Sequence Detection System (Applied Biosystems, Foster City, CA, USA) was used for qPCR analysis, following the manufacturer’s instructions. To analyze ERα target gene expression, the PrimePCR Estrogen receptor signaling (SAB Target List) H96 panel (Bio-Rad Laboratories, Hercules, CA, USA) was employed for RT-qPCR-based gene array analysis, as per the manufacturer’s instructions. Gene expression data were normalized to the levels of GAPDH mRNA present in the array. Genes were considered affected if their fold induction was above 1.5 or below 0.7 compared to the control sample.

### Affimetrix analysis

Total RNA was extracted using RNeasy kit (Qiagen), according to manufacturer’s protocol, and was quantified using a NanoDrop 2000 system (Thermo Scientific). A GeneChip Pico Reagent Kit (Affymetrix) was used to amplify 5 ng of total RNA, according to the manufacturer’s protocol. Quality control of the RNA samples was performed using an Agilent Bioanalyzer 2100 system (Agilent Technologies). Gene expression profiling was performed using the Affymetrix GeneChip® Human Clariom S Array (Thermo Fisher Scientific), including more than 210,000 distinct probes representative of 21,448 annotated genes (Genome Reference Consortium Human Build 38 (GRCh38); https://www.ncbi.nlm.nih.gov/datasets/genome/GCF_000001405.26/). RNA samples were amplified, fragmented, and labeled for array hybridization according to manufacturer’s instruction. Samples were then hybridized overnight, washed, stained, and scanned using the Affymetrix GeneChip Hybridization Oven 640, Fluidic Station 450 and Scanner 3000 7G (Thermo Fisher Scientific), to generate raw data files (.CEL files). Quality control and normalization of Affymetrix .CEL files were performed using the TAC software (v4.0; Thermo Fisher Scientific), by performing the “Gene level SST-RMA” summarization method with human genome version hg38 (https://www.ncbi.nlm.nih.gov/assembly/GCF_000001405.26/). Gene expression data were log2 transformed before analyses. Class comparison analysis for identifying differentially regulated genes was performed using TAC software by selecting a fold-change (FC) of |2| and FDR adjusted p-value (Benjamini-Hochberg Step-Up FDR-controlling Procedure) ≤ 0.05 as cutoff.

### 5-ethynyl-2’-deoxyuridine (EdU) Incorporation Assay

The cell medium was supplemented with 5-ethynyl-2’-deoxyuridine (EdU) during the last 30 minutes of cell growth. After the EdU incubation, the cells were fixed and permeabilized. The EdU assay was performed using the Click-iT™ EdU Cell Proliferation Kit for Imaging, Alexa Fluor™ 488 dye, following the manufacturer’s instructions. Fluorescence was measured directly in 96-well plates, with each sample being repeated at least in triplicate. The measurements were performed using a Tecan Spark Reader.

### Statistical Analysis

Statistical analysis was conducted using InStat version 8 software system (GraphPad Software Inc., San Diego, CA). Densitometric analyses were carried out using Image J freeware software, where the band intensity of the protein of interest was quantified relative to the loading control band (vinculin) intensity. The p-values and the specific statistical test used (either Student t-test or ANOVA Test) are provided in the figure captions.

## List of Abbreviations

AI: Aromatase inhibitors
AKT: V-Akt Murine Thymoma Viral Oncogene Homolog 1
ALK: Anaplastic lymphoma kinase
AP: AP26113
ATR: Ataxia Telangiectasia And Rad3-Related Protein
AURKA: Aurora kinase A
AURKB: Aurora kinase B
Baf: Bafilomycin A1
BC: breast cancer
BDNF: Brain Derived Neurotrophic Factor
BRCA1: Breast cancer type 1 susceptibility protein
BUB1: Mitotic Checkpoint Serine/Threonine-Protein Kinase BUB1
BUB1B: BUB1 Mitotic Checkpoint Serine/Threonine Kinase B
CAMK2D: Calcium/Calmodulin Dependent Protein Kinase II Delta
CDC7: Cell Division Cycle 7
CDK1: cyclin-dependent kinase 1
CDK2: cyclin-dependent kinase 2
CHK1: Checkpoint Kinase 1
CHX: Cycloheximide
CycD1: cyclin D1
DCLK1: Doublecortin Like Kinase 1
DMEM: Dulbecco’s Midified Eagle Medium
DYRK1B: Dual Specificity Tyrosine Phosphorylation Regulated Kinase 1B
E2: 17β-estradiol
EC50: Effective concentration 50
EdU: 5-ethynyl-2’-deoxyuridine
ERE: estrogen responsive element
Erlo: Erlotinib
ERα: estrogen receptor α
ET: Endocrine therapy
FDA: Food and Drug Administration
FOXA1: Forkhead Box A1
GART: Phosphoribosylglycinamide Formyltransferase, Phosphoribosylglycinamide Synthetase, Phosphoribosylaminoimidazole Synthetase
Gef: Geftinib
GSG2: Histone H3 Associated Protein Kinase
HER2: Human Epidermal Growth Factor Receptor 2
IC50: Inhibitory concentration 50
IDC: Invasive ductal carcinoma
IGF-1R: Insulin-like growth factor 1 receptor
Kd: Dissociation constant
Lapa: Lapatinib
LumA: Luminal A
LumB: Luminal B
MASTL: Microtubule Associated Serine/Threonine Kinase Like
MBC: Metastatic breast cancer
MELK: Maternal Embryonic Leucine Zipper Kinase
MELKin: MELK-8a – MELK inhibitor
mRNA: Messenger ribonucleic acid
NLuc: Nanoluciferase
p62^SQSTM^: protein 62/sequestrosome
PBK: PDZ Binding Kinase
PLK: Polo-like kinase
PLK4: Polo-like kinase 4
PR: Progesterone Receptor
pS2: presenelin2
RARA: retinoic acid receptor alpha
RFS: relapse free survival
Tam: 4OH-tamoxifen
Tel: Telaprevir
TTK: Phosphotyrosine Picked Threonine-Protein Kinase
UPS: Ubiquitin proteasome system
VRK1: VRK Serine/Threonine Kinase 1
YY: Buffer: Yoss Yarden Buffer

## Acknowledgements

The research leading to these results has received funding from AIRC under IG 2018 - ID. 21325 project – P.I. Acconcia Filippo. This study was also supported by grants from Ateneo Roma Tre to FA. The Grant of Excellence Departments 2023-2027, MIUR (ARTICOLO 1, COMMI 314 – 337 LEGGE 232/2016) to the Department of Science, University Roma TRE is also gratefully acknowledged. The authors are grateful to Prof. Simak Ali, University of London Imperial College for the gift of the MCF-7 Y537S cells.

## Authors’ contributions

SB performed all the experiments regarding MELK and some regarding ALK. SP performed almost all the experiments regarding ALK. FB performed Affymetrix analyses. MC performed experiments in 3D model system of breast cancer cells. FA performed the in-silico evaluations, analyzed the data, conceived the experiments, wrote the paper. All authors, which contributed to manuscript revision and editing, read and approved the final manuscript.

## Data availability Statement

All the original Western blots with replicates of the experiments are available as a separate file uploaded together with this work. All the Kaplan-Meier curves were retrieved by the Kaplan-Meier Plotter database and given in supplementary table 5 as downloaded by the website (https://kmplot.com/analysis/) ^28^. All the datasets used to generate figure 1 were downloaded by the Broad Institute through the DepMap portal (https://depmap.org/portal) and are available in supplementary table 1 and table 2. Datasets used to generate figure 2A and 2B are given in supplementary table 3. Data used to generate figure 2C, 2D and 2E are given in supplementary table 4. The results of the esiRNA and the three ALK, AURKA/AURKB inhibitor screenings in the seven breast cancer cell lines for measuring the ERα levels as well as those for growth curve analyses, which were produced and analyzed during the current study, are available from the corresponding author upon reasonable request.

## References

1 Morganti, S. C., G. Moving beyond endocrine therapy for luminal metastatic breast cancer in the precision medicine era: looking for new targets. EXPERT REVIEW OF PRECISION MEDICINE AND DRUG DEVELOPMENT, doi:10.1080/23808993.2020.1720508 (2020).

2 Parsons, J. & Francavilla, C. ’Omics Approaches to Explore the Breast Cancer Landscape. Front Cell Dev Biol 7, 395, doi:10.3389/fcell.2019.00395 (2019).

3 Johansson, H. J. et al. Breast cancer quantitative proteome and proteogenomic landscape. Nature communications 10, 1600, doi:10.1038/s41467-019-09018-y (2019).

4 Tsang, J. Y. S. & Tse, G. M. Molecular Classification of Breast Cancer. Adv Anat Pathol 27, 27–35, doi:10.1097/PAP.0000000000000232 (2020).

5 Lumachi, F. et al. Endocrine therapy of breast cancer. Curr Med Chem 18, 513–522, doi:not available. PMID: 21143113. (2011).

6 Acconcia, F. Evaluation of the Sensitivity of Breast Cancer Cell Lines to Cardiac Glycosides Unveils ATP1B3 as a Possible Biomarker for the Personalized Treatment of ERalpha Expressing Breast Cancers. International journal of molecular sciences 23, doi:10.3390/ijms231911102 (2022).

7 Cipolletti, M. et al. A new anti-estrogen discovery platform identifies FDA-approved imidazole anti-fungal drugs as bioactive compounds against ERα expressing breast cancer cells. International journal of molecular sciences 22, doi:10.3390/ijms22062915 (2021).

8 Busonero, C., Leone, S., Bartoloni, S. & Acconcia, F. Strategies to degrade estrogen receptor alpha in primary and ESR1 mutant-expressing metastatic breast cancer. Mol Cell Endocrinol 480, 107–121, doi:10.1016/j.mce.2018.10.020 (2019).

9 Busonero, C., Leone, S. & Acconcia, F. Emetine induces estrogen receptor alpha degradation and prevents 17beta-estradiol-induced breast cancer cell proliferation. Cellular oncology, doi:10.1007/s13402-017-0322-z (2017).

10 Busonero, C. et al. Ouabain and Digoxin Activate the Proteasome and the Degradation of the ERα in Cells Modeling Primary and Metastatic Breast Cancer. Cancers (Basel) 12, doi:10.3390/cancers12123840 (2020).

11 Busonero, C., Leone, S., Klemm, C. & Acconcia, F. A functional drug re-purposing screening identifies carfilzomib as a drug preventing 17beta-estradiol: ERalpha signaling and cell proliferation in breast cancer cells. Mol Cell Endocrinol 460, 229–237, doi:10.1016/j.mce.2017.07.027 (2018).

12 Busonero, C. L., S.; Bianchi, F.; Acconcia, F. In silico screening for ERα downmodulators identifies thioridazine as an anti-proliferative agent in primary, 4OH-tamoxifen-resistant and Y537S ERα-expressing breast cancer cells. Cellular oncology, doi:10.1007/s13402-018-0400-x (2018).

13 Cipolletti, M., Leone, S., Bartoloni, S., Busonero, C. & Acconcia, F. Real-time measurement of E2: ERalpha transcriptional activity in living cells. J Cell Physiol, doi:10.1002/jcp.29565 (2020).

14 Leone, S., Busonero, C. & Acconcia, F. A high throughput method to study the physiology of E2:ERalpha signaling in breast cancer cells. J Cell Physiol 233, 3713–3722, doi:10.1002/jcp.26251 (2018).

15 Acconcia, F. et al. The extra-nuclear interactome of the estrogen receptors: implications for physiological functions. Mol Cell Endocrinol 538, 111452, doi:10.1016/j.mce.2021.111452 (2021).

16 Cipolletti, M., Leone, S., Bartoloni, S. & Acconcia, F. A functional genetic screen for metabolic proteins unveils GART and the de novo purine biosynthetic pathway as novel targets for the treatment of luminal A ERalpha expressing primary and metastatic invasive ductal carcinoma. Front Endocrinol (Lausanne) 14, 1129162, doi:10.3389/fendo.2023.1129162 (2023).

17 Pescatori, S. et al. Clinically relevant CHK1 inhibitors abrogate wild-type and Y537S mutant ERα expression and proliferation in luminal primary and metastatic breast cancer cells. Journal of Experimental & Clinical Cancer Research 41, 27, doi:10.1186/s13046-022-02360-y (2022).

18 Neve, R. M. et al. A collection of breast cancer cell lines for the study of functionally distinct cancer subtypes. Cancer cell 10, 515–527, doi:10.1016/j.ccr.2006.10.008 (2006).

19 Dai, X., Cheng, H., Bai, Z. & Li, J. Breast Cancer Cell Line Classification and Its Relevance with Breast Tumor Subtyping. J Cancer 8, 3131–3141, doi:10.7150/jca.18457 (2017).

20 Techer, H. & Pasero, P. The Replication Stress Response on a Narrow Path Between Genomic Instability and Inflammation. Front Cell Dev Biol 9, 702584, doi:10.3389/fcell.2021.702584 (2021).

21 Bhola, N. E. et al. Kinome-wide functional screen identifies role of PLK1 in hormone-independent, ER-positive breast cancer. Cancer Res 75, 405–414, doi:10.1158/0008-5472.CAN-14-2475 (2015).

22 Bhola, N. E. et al. Correction: Kinome-wide Functional Screen Identifies Role of PLK1 in Hormone-Independent, ER-Positive Breast Cancer. Cancer Res 79, 876, doi:10.1158/0008-5472.CAN-18-4088 (2019).

23 Bartoloni, S., Leone, S. & Acconcia, F. Unexpected Impact of a Hepatitis C Virus Inhibitor on 17beta-Estradiol Signaling in Breast Cancer. International journal of molecular sciences 21, doi:10.3390/ijms21103418 (2020).

24 Bartoloni, S., Leone, S., Pescatori, S., Cipolletti, M. & Acconcia, F. The antiviral drug telaprevir induces cell death by reducing FOXA1 expression in estrogen receptor alpha (ERalpha)-positive breast cancer cells. Mol Oncol 16, 3568–3584, doi:10.1002/1878-0261.13303 (2022).

25 Harrod, A. et al. Genomic modelling of the ESR1 Y537S mutation for evaluating function and new therapeutic approaches for metastatic breast cancer. Oncogene 36, 2286–2296, doi:10.1038/onc.2016.382 (2017).

26 Harrod, A. et al. Genome engineering for estrogen receptor mutations reveals differential responses to anti-estrogens and new prognostic gene signatures for breast cancer. Oncogene 41, 4905–4915, doi:10.1038/s41388-022-02483-8 (2022).

27 Finetti, P. et al. Sixteen-kinase gene expression identifies luminal breast cancers with poor prognosis. Cancer Res 68, 767–776, doi:10.1158/0008-5472.CAN-07-5516 (2008).

28 Lanczky, A. & Gyorffy, B. Web-Based Survival Analysis Tool Tailored for Medical Research (KMplot): Development and Implementation. J Med Internet Res 23, e27633, doi:10.2196/27633 (2021).

29 Toure, B. B. et al. Toward the Validation of Maternal Embryonic Leucine Zipper Kinase: Discovery, Optimization of Highly Potent and Selective Inhibitors, and Preliminary Biology Insight. Journal of medicinal chemistry 59, 4711–4723, doi:10.1021/acs.jmedchem.6b00052 (2016).

30 Klionsky, D. J. et al. Guidelines for the use and interpretation of assays for monitoring autophagy. Autophagy 8, 445–544, doi:10.4161/auto.19496 (2012).

31 Creighton, C. J. The molecular profile of luminal B breast cancer. Biologics 6, 289–297, doi:10.2147/BTT.S29923 (2012).

32 Ali, S., Metzger, D., Bornert, J. M. & Chambon, P. Modulation of transcriptional activation by ligand-dependent phosphorylation of the human oestrogen receptor A/B region. EMBO J 12, 1153–1160, doi:not available. PMID: 8458328. (1993).

33 Langhans, S. A. Three-Dimensional in Vitro Cell Culture Models in Drug Discovery and Drug Repositioning. Front Pharmacol 9, 6, doi:10.3389/fphar.2018.00006 (2018).

34 Bartlett, J. M. S. & Parelukar, W. Breast cancers are rare diseases-and must be treated as such. NPJ breast cancer 3, 11, doi:10.1038/s41523-017-0013-y (2017).

35 Duong-Ly, K. C. & Peterson, J. R. The human kinome and kinase inhibition. Curr Protoc Pharmacol Chapter 2, Unit2 9, doi:10.1002/0471141755.ph0209s60 (2013).

36 Totta, P., Busonero, C., Leone, S., Marino, M. & Acconcia, F. Dynamin II is required for 17beta-estradiol signaling and autophagy-based ERalpha degradation. Scientific reports 6, 23727, doi:10.1038/srep23727 (2016).

37 Korolchuk, V. I., Mansilla, A., Menzies, F. M. & Rubinsztein, D. C. Autophagy inhibition compromises degradation of ubiquitin-proteasome pathway substrates. Mol Cell 33, 517–527, doi:10.1016/j.molcel.2009.01.021 (2009).

38 Liu, W. J. et al. p62 links the autophagy pathway and the ubiqutin-proteasome system upon ubiquitinated protein degradation. Cell Mol Biol Lett 21, 29, doi:10.1186/s11658-016-0031-z (2016).

39 Laios, I. et al. Role of the proteasome in the regulation of estrogen receptor alpha turnover and function in MCF-7 breast carcinoma cells. J Steroid Biochem Mol Biol 94, 347–359, doi:10.1016/j.jsbmb.2005.02.005 (2005).

40 Gettinger, S. N. et al. Activity and safety of brigatinib in ALK-rearranged non-small-cell lung cancer and other malignancies: a single-arm, open-label, phase 1/2 trial. Lancet Oncol 17, 1683–1696, doi:10.1016/S1470-2045(16)30392-8 (2016).

41 Cipolletti, M., Pescatori, S. & Acconcia, F. Real-time challenging of ERα Y537S mutant transcriptional activity in living cells. Endocrines 2, 54–64, doi:10.3390/endocrines2010006 (2021).

42 Totta, P., Pesiri, V., Enari, M., Marino, M. & Acconcia, F. Clathrin Heavy Chain Interacts With Estrogen Receptor alpha and Modulates 17beta-Estradiol Signaling. Mol Endocrinol 29, 739–755, doi:10.1210/me.2014-1385 (2015).

43 Pesiri, V., Totta, P., Marino, M. & Acconcia, F. Ubiquitin-activating enzyme is necessary for 17beta-estradiol-induced breast cancer cell proliferation and migration. IUBMB Life 66, 578–585, doi:10.1002/iub.1296 (2014).

