## Supplementary material for "Selective Impact of ALK and MELK Inhibition on ERα Stability and Cell Proliferation in Cell Lines Representing Distinct Molecular Phenotypes of Breast Cancer": Caption of Supplementary Tables.docx

**Supplementary Table 1**

Mean and p-Values of the sensitivity data of each cell line to each drug as downloaded by DepMap. The file also includes the lists of the identified drugs and the identified kinase inhibitors.

**Supplementary Table 2**

The file includes the sensitivities of each listed cell line both to the antiproliferative action and to the ERα degradation of ALK and AURKA/AURKB inhibitors.

**Supplementary Table 3**

The file includes the list of the downmodulated kinases in the cell lines used for telaprevir Affymetrix analyses and also those kinases that are in common with that identified in Finetti et al., 2008.

**Supplementary Table 4**

The file includes the sensitivities of each listed cell line both to the esiRNA treatment against the different to ERα degradation.

**Supplementary Table 5**

The file includes the original data used for Kaplan-Meier analyses as downloaded by the kmplotter database.
