## Supplementary material for "Selective Impact of ALK and MELK Inhibition on ERα Stability and Cell Proliferation in Cell Lines Representing Distinct Molecular Phenotypes of Breast Cancer": graphical abstract.pptx

### Slide 1
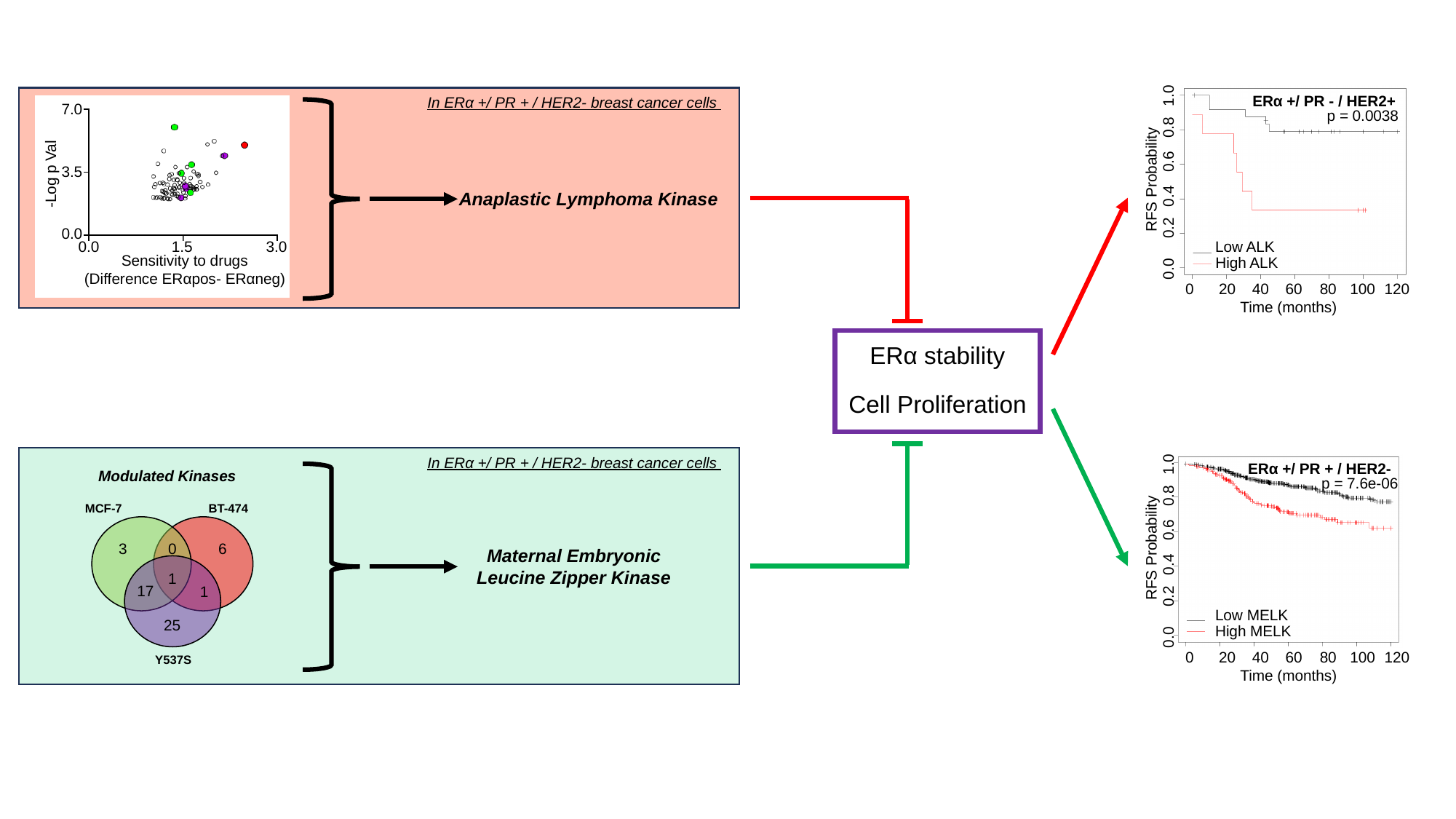

1.0
ERα +/ PR - / HER2+
p = 0.0038
0.8
0.6
RFS Probability
0.4
0.2
Low ALK
High ALK
0.0
0
20
40
60
80
100
120
Time (months)
In ERα +/ PR + / HER2- breast cancer cells
7.0
3.5
-Log p Val
0.0
0.0
1.5
3.0
Sensitivity to drugs
(Difference ERαpos- ERαneg)
Anaplastic Lymphoma Kinase
ERα stability
Cell Proliferation
1.0
ERα +/ PR + / HER2-
p = 7.6e-06
0.8
0.6
RFS Probability
0.4
0.2
Low MELK
High MELK
0.0
0
20
40
60
80
100
120
Time (months)
In ERα +/ PR + / HER2- breast cancer cells
Modulated Kinases
MCF-7
BT-474
3
0
6
1
17
1
25
Y537S
Maternal Embryonic Leucine Zipper Kinase
