## Supplementary material for "Selective Impact of ALK and MELK Inhibition on ERα Stability and Cell Proliferation in Cell Lines Representing Distinct Molecular Phenotypes of Breast Cancer": Supplementary Figure Captions.docx

**Figure 1. Correlation between two ALK inhibitors.**

Linear regression and Spearman Correlation values for the effective concentration 50 (EC_50_) for inhibitor-induced reduction in ERα intracellular levels between the ALK inhibitors AZD3436 – AZD and and AP26113 – AP in all the seven BC cell lines used in the study. Main panels display the correlation coefficient (r) and p-values.
