## Supplementary figures and images for "Selective Impact of ALK and MELK Inhibition on ERα Stability and Cell Proliferation in Cell Lines Representing Distinct Molecular Phenotypes of Breast Cancer"

### Supplementary Figures.pptx

#### Slide 1
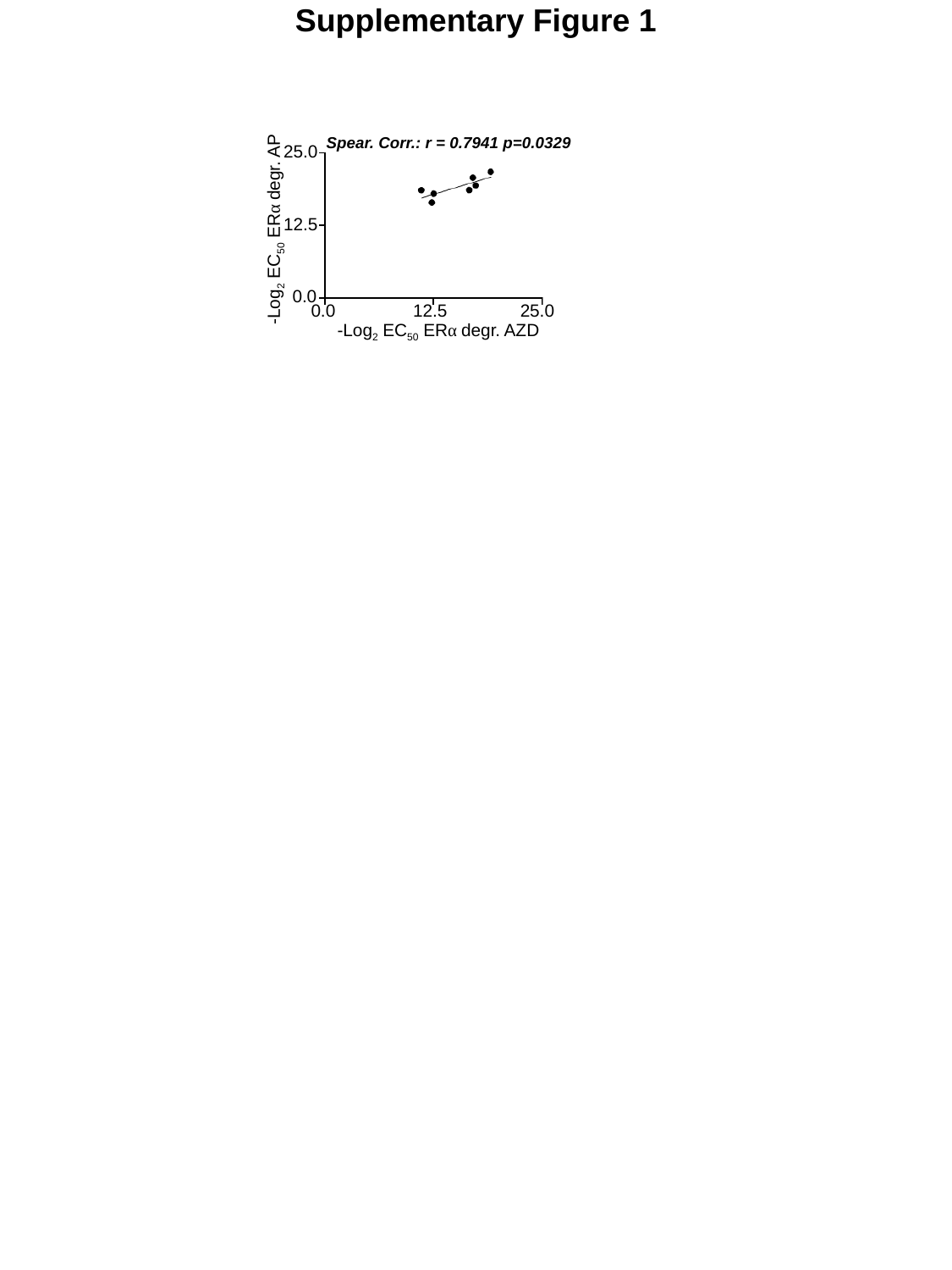

Supplementary Figure 1
Spear. Corr.: r = 0.7941 p=0.0329
25.0
12.5
-Log_2_ EC_50_ ERα degr. AP
0.0
0.0
12.5
25.0
-Log_2_ EC_50_ ERα degr. AZD
